# Structure-Guided Loop Grafting Improves Expression and Stability of Influenza Neuraminidase for Vaccine Development

**DOI:** 10.1101/2024.10.11.617814

**Authors:** Pramila Rijal, Leiyan Wei, Guido C Paesen, David I Stuart, Mark R Howarth, Kuan-Ying A Huang, Thomas A Bowden, Alain RM Townsend

## Abstract

Influenza virus neuraminidase is a crucial target for protective antibodies, yet the development of recombinant neuraminidase protein as a vaccine has been held back by instability and variable expression. We have taken a pragmatic approach to improving expression and stability of neuraminidase by grafting antigenic surface loops from low-expressing neuraminidase proteins onto the scaffold of high-expressing counterparts. The resulting hybrid proteins retained the antigenic properties of the loop donor while benefiting from the high-yield expression, stability, and tetrameric structure of the loop recipient. These hybrid proteins were recognised by a broad set of human monoclonal antibodies elicited by influenza infection or vaccination, with X-ray structures validating the accurate structural conformation of the grafted loops and the enzymatic cavity. Immunisation of mice with neuraminidase hybrids induced inhibitory antibodies to the loop donor and protected against lethal influenza challenge. This pragmatic technique offers a robust solution for improving the expression and stability of influenza neuraminidase proteins for vaccine development.

**Graphical Abstract:** 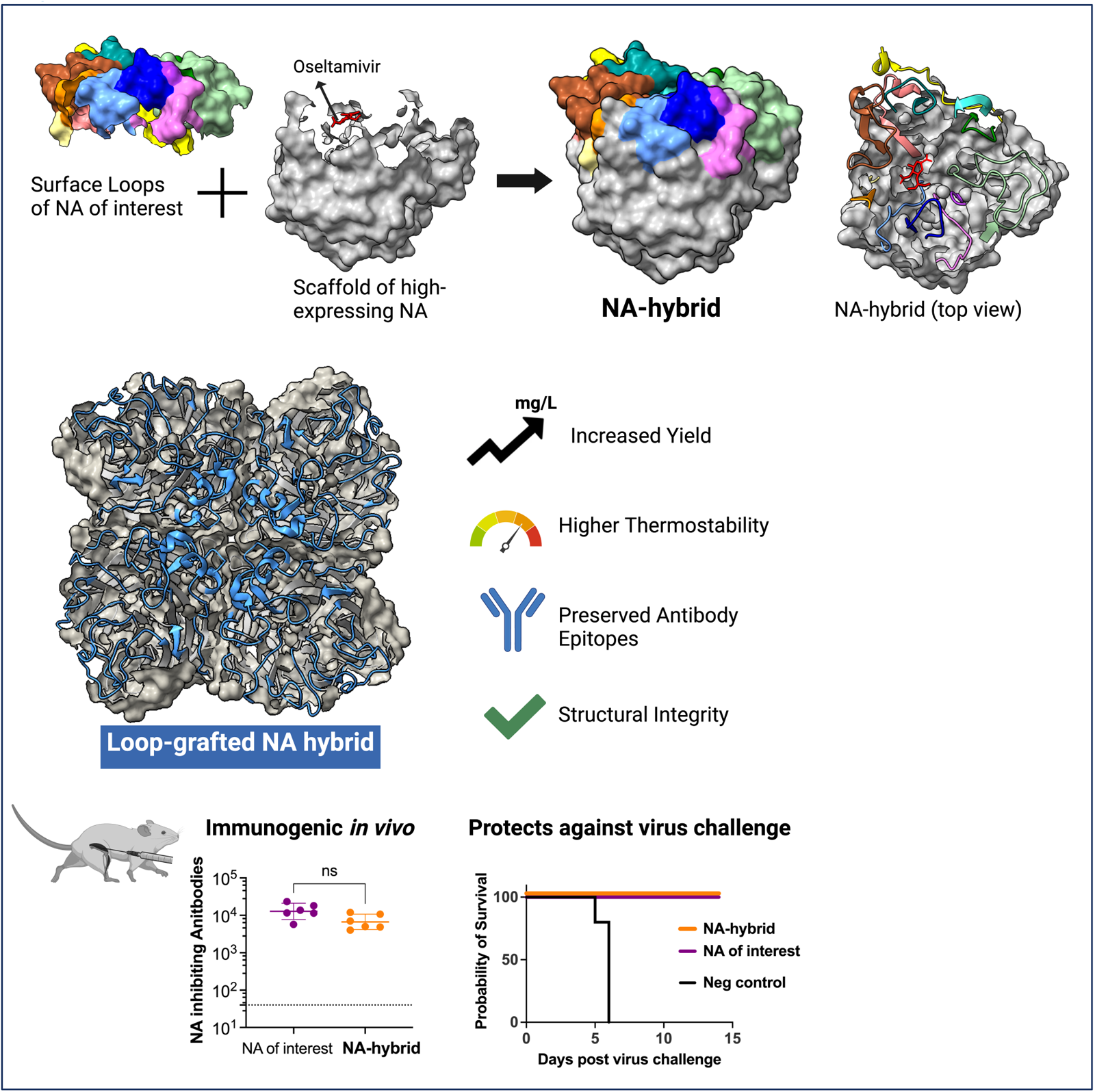

## Introduction

Influenza virus neuraminidase (NA) is an important target for protective antibodies and therapeutic drugs against influenza viruses. Currently, influenza subunit vaccines focus on haemagglutinin and contain minimal amounts of NA, despite emerging evidence from multiple studies indicating NA’s importance as an independent protective antigen ^1^ (reviewed in ^2–6^). However, recombinant NA protein expression is often bedevilled by low yield and poor stability, with substantial variation in NA protein yields across different virus strains (Supplementary Table 1). With the ExpiCHO transient expression system (Thermo Fisher), we have found that yields of the N1 subtype NA varied from undetectable (H1N1/2019) to a maximum of 380 mg per L of culture (H5N1/2021).

We have applied structural and epitope information to devise a solution to improve the expression and stability of our recombinant NAs. NA is a mushroom shaped type II homotetrameric protein ^7^ reviewed in ^2^). The polypeptide chain of the NA monomeric head folds into six, topologically identical, four-stranded, antiparallel β-sheets which are arranged like the blades of a propeller ^7^. Each of the six β-sheet blades, numbered B1-B6, is composed of four anti-parallel β-strands in a W-shape numbered S1-S4. The β-strands in the “W” are connected at the top and bottom by loops (Fig. 1a). The loop between the fourth strand of the preceding beta sheet and the first strand of the following sheet (L01), and the loop between the second and third strands (L23) form most of the top surface and the active site of the NA and were delineated in the first crystal structure of the N2 NA ^7,8^ and in subsequent structures (reviewed in ^2^). As there are six β-sheets in each NA monomer, there are six L01 and six L23 loops that together form most of the top surface of each identical NA monomer with a final contribution from the C-terminal domain of ∼17 amino acids (Fig. 1a).

**Fig. 1:**
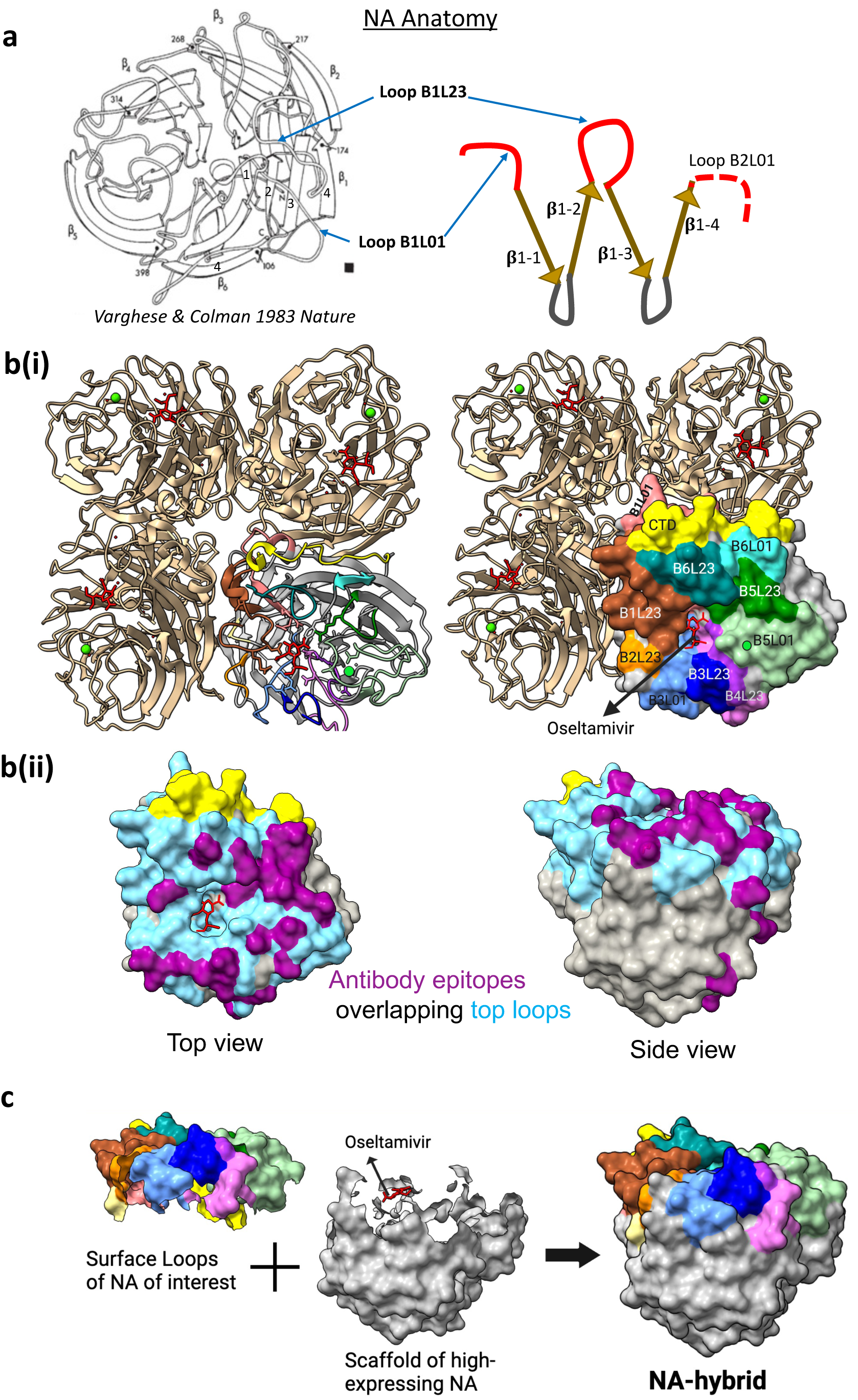
Surface antigenic loops transfer method to design NA-hybrid protein. **a) NA Anatomy.** Each monomer of the tetrameric NA head is composed of a six-bladed propeller-like structure. Each blade unit consists of a β-sheet composed of four β-strands connected by loops arranged in a W-shape. The loop preceding β-strand 1 is termed Loop01 and the loop connecting β-strands 2 and 3 is termed Loop23. The twelve 01 and 23 loops in each monomer point up and surround and contribute to the enzyme active site. **b) Antigenic surface formation. (i)** The key antigenic surface on the top of each NA monomer is formed by the twelve 01 and 23 loops from the six β-sheet units and the C-terminal domain. The twelve loops and the C-terminal domain in one monomer are coloured to show the top surface surrounding the active site. The remaining four Beta strands and the Loops 12 and 34 we refer to as the “scaffold”. Oseltamivir is shown in red in the sialic acid receptor binding site. A calcium ion at its binding site is shown as a green circle. (**ii)** Twelve L12 and L23 loops are coloured cyan blue and antibody epitopes as purple (L12 & L23 loops represent 90% of antibody epitopes - refer to Supplementary Figure 1 for epitope mapping on the aligned sequences). Similar to b(i), oseltamivir and C-terminal domain are shown for orientation. **c) Hybrid NA design.** Loops 01 and 23 and the C-terminal domain, and the scaffold are shown separately. Oseltamivir and the binding residues are included for reference. The surface antigenic loops 01 and 23 of an NA of interest were transferred on to the scaffold of a high-expressing NA candidate to improve the protein yield and stability. Figures were generated with PDB 4B7J using UCSF ChimeraX^22^.

The L01 and L23 loops encircle the cavity forming the NA enzyme active site that binds sialic acid. NA cleaves off host sialic acids, aiding the release of newly formed virions from infected cells ^9^. While the loop residues contributing to the active site of NA are completely conserved, the remaining sequence of these loops can be highly variable. Many antibodies that inhibit enzyme activity and are protective in vivo bind to these loops (^8,10–13^, reviewed in ^14^, Supplementary Fig. 1, 2). Multiple studies have also shown that the majority of, but not all (Supplementary Fig. 1), monoclonal antibodies (mAbs) to NA select resistant viruses with amino acid replacements in the L01 and L23 loops. Analysis of the evolution of NAs has shown that the great majority of sequence change over time occurs at the surface exposed residues of NA, which include the residues of the L01 and L23 loops ^8,15^.

We assumed that the interactions between monomers in the NA tetramer are likely to make a major contribution to the stability and efficiency of expression of recombinant tetrameric NA *in vitro.* From known crystal structures Ellis *et al*. ^16^ identified forty-four residues that may contribute to these contacts in the N1 protein, ten of which occurred in loops L01 and L23 (as defined by Varghese *et al*. ^7^). In their final design of the stabilised sNAp-155 N1 2009 NA they replaced ten of these forty-four amino acids, of which only two residues (conserved in N1 NA) were in the surface loops B2L01 and B2L23. Thus, the majority of intermonomer contacts are *not* in the L01 and L23 surface loops. In addition, the tertiary structure of NA is highly conserved. Within a NA subtype the Cα atoms of the NA head (including surface loops), are superimposable with average root mean square deviation (RMSD) between the Cα atoms of less than 0.37 Å for N1 ^17,18^. From these observations we reasoned that it might be possible to combine the antigenic features of a poorly expressed NA with the expression and stability properties of a high expressing NA by grafting the L01 and L23 loops from the former onto the structural “scaffold” of the latter, particularly within an NA subtype.

## Results

### Design of loop-grafted NA hybrid proteins

NA proteins were expressed in a transient mammalian ExpiCHO expression system using a gene construct comprising the N-terminus signal sequence, purification tags, SpyTag, a tetrabrachion tetramerization domain ^19^ and the NA head ectodomain (Supplementary Fig. 3a, see Methods). The protein was purified from clarified supernatants and analysed using size-exclusion chromatography, SDS-PAGE with BS3 cross linking, thermal unfolding, and enzymatic assays.

Using the PDB 1NN2 (H2N2 A/Tokyo/3/1967) structure and the top surface loop annotations by Varghese *et al.* ^7^ as a guide, we annotated the twelve L01 and L23 loops of the N1 neuraminidases (Fig. 1 a,b, Fig. 2, Supplementary Fig. 4, Supplementary Table 2). We transferred 12 dissimilar residues within the loops of N1/09 and 16 dissimilar residues of N1/19 to the H5N1/2021 scaffold (mSN1) to create N1/09 and N1/19 hybrids (Fig. 2a,b). From now on the protein containing the N1/09Loops on the mS scaffold and N1/19 Loops on the mS scaffold will be referred to as the N1/09 hybrid and N1/19 hybrid, respectively. The C-terminal domain (CTD) of the final ∼17 amino acids of the N1 protein also forms part of the top surface. However, in these N1 NAs the CTD was conserved.

**Fig 2:**
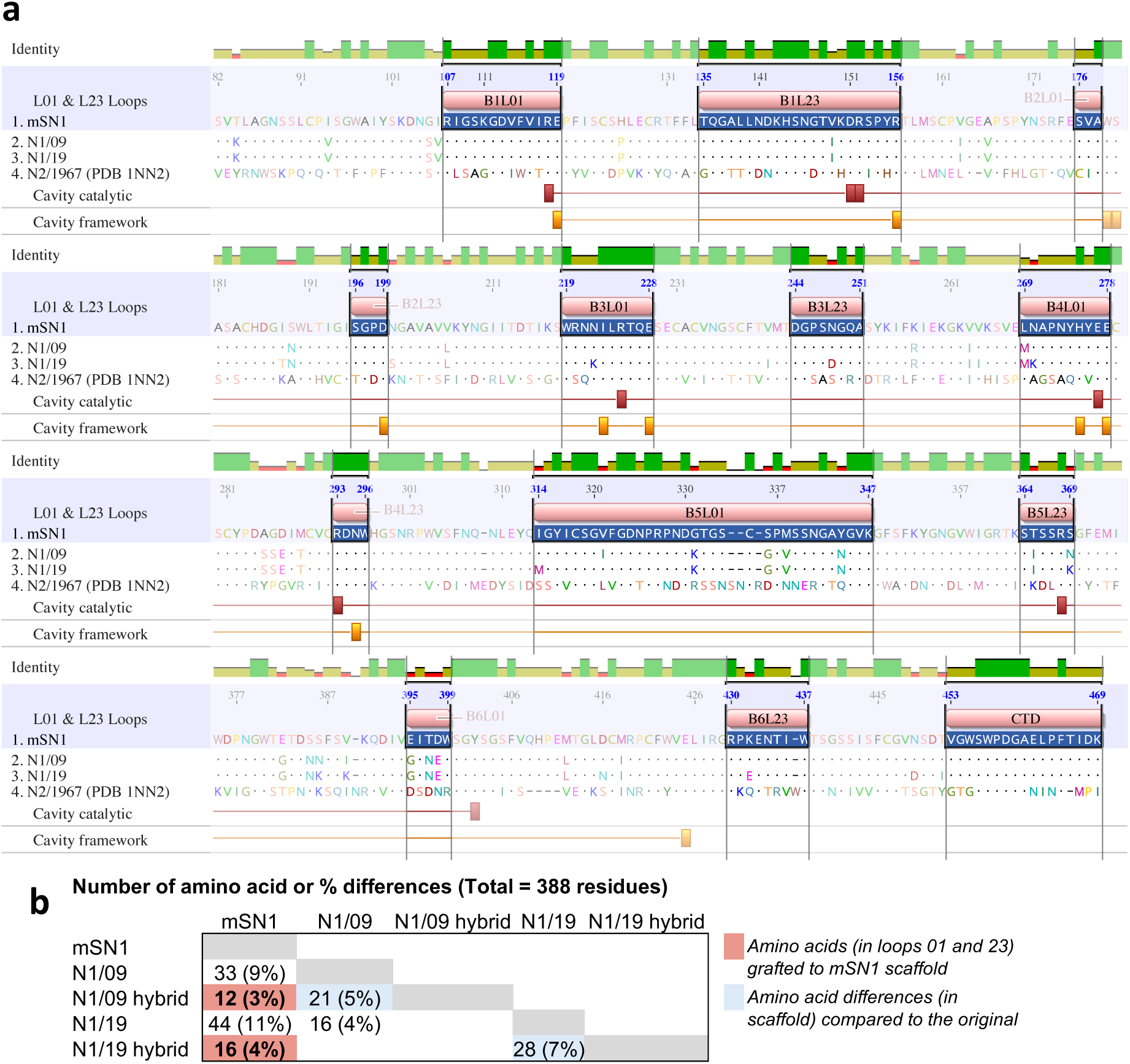
NA protein sequence alignment highlighting surface loops and active site. **a)** Protein sequence alignment of the scaffold of N1 from H5N1 A/mute swan/England/053054/2021 (mS), and loop donors H1N1 A/California/7/2009 (N1/09) and A/Wisconsin/588/2019 (N1/19) neuraminidases. The N2 sequence H2N2 A/Tokyo/3/1967 from Varghese et. al (1983, 1991), that was used to define the loops (PDB 1NN2) is also included. mSN1 is used as a reference sequence and identical residues are shown as dots. The sequence conservation is shown by green bars. The numbering here is based on N1 numbering as used in the annual Crick Reports (https://www.crick.ac.uk/sites/default/files/2022-10/Crick%20report%20Sep2022%20for%20SH2023_to%20post.pdf). Loops 01 and 23 that form the top antigenic surface are highlighted. Loops annotation is based on Varghese et al. (see Supplementary Table 2 and Supplementary Figure 4 for detailed information). Residues that form the catalytic site (8 residues) and conserved framework residues for the catalytic site (11 residues) of the enzymatic cavity are annotated with red and orange bars respectively. These residues are highly conserved between NA subtypes and the majority are part of the surface loops - 7/8 catalytic residues and 8/11 catalytic site framework residues. Two catalytic site framework residues are at the edge of loop B2L01. The figure was generated using Geneious Prime. **b)** Number of amino acid differences for the N1/09 and N1/19 loop donors and the mS N1 loop recipient are shown. Twelve dissimilar residues within Loops 01 and 23 of N1/09 were transferred to mSN1 scaffold to form N1/09 hybrid. Similarly, 16 dissimilar residues within Loops 01 and 23 of N1/19 were transferred to mS N1 scaffold to form N1/19 hybrid.

### N1/09 and N1/19 hybrid proteins expressed at higher level as stable tetramers

We found that H5N1 A/mute swan/England/053054/2021 NA (mSN1) expressed at ∼380 mg/L as a tetramer with minimal aggregation, exhibiting high enzymatic activity and a high melting temperature (Fig. 3a). However, H1N1 A/California/07/2009 NA (N1/09) expressed at ∼15 mg/L with a tendency to form aggregates, and H1N1 A/Wisconsin/588/2019 NA (N1/19) had a variable expression from undetectable to low (performed twice).

**Fig 3:**
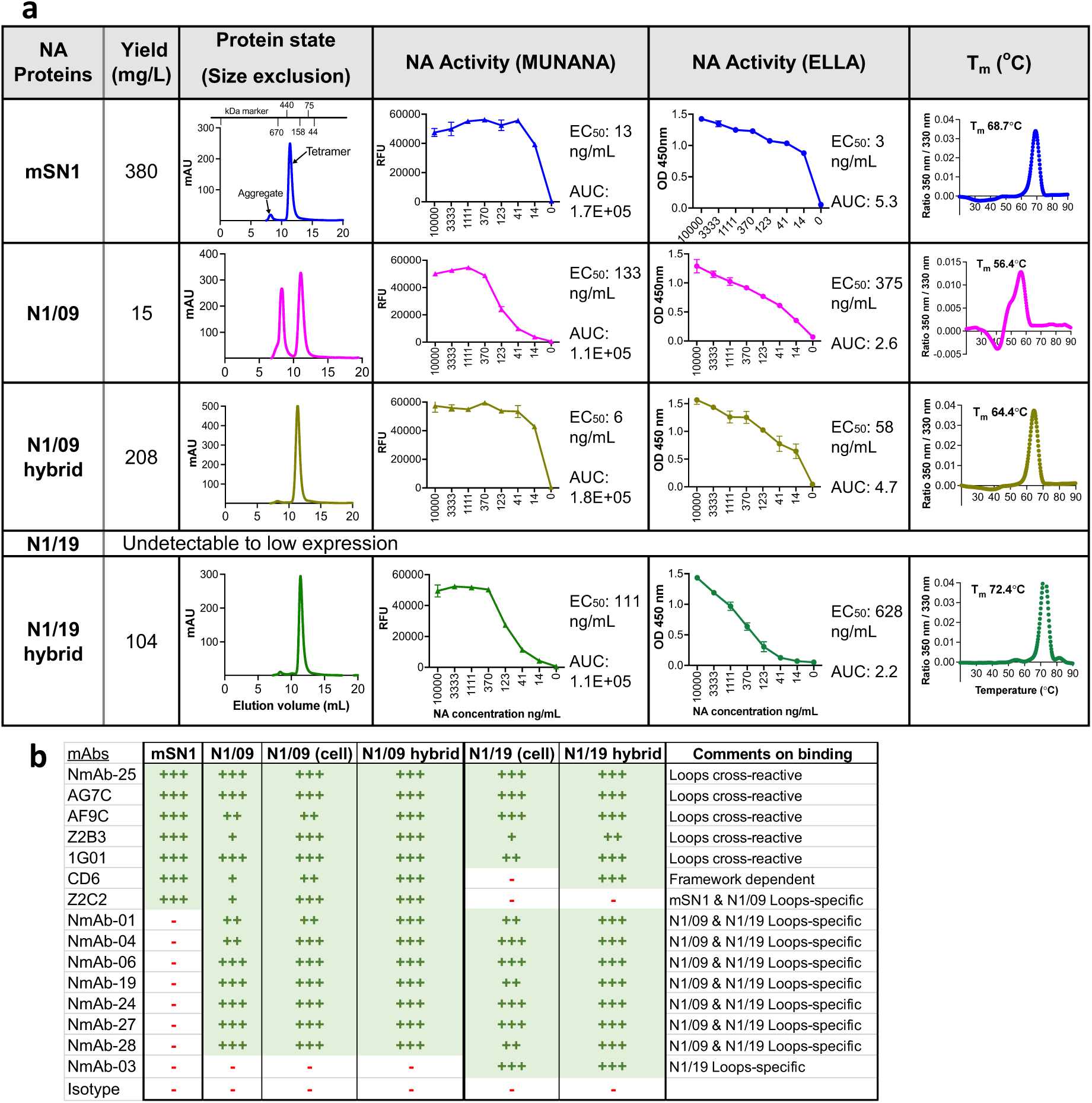
Characteristics of NA hybrid proteins including epitope specificity. **a) Characteristics of NA proteins.** NA proteins were expressed in a transient mammalian ExpiCHO expression system. Gene constructs including affinity purification tags, SpyTag and an artificial tetramerisation domain tetrabrachion at N-terminus were cloned in the pcDNA3.1-vector. Proteins were purified using the 6His tag and Nickel-sepharose HisTrap purification columns. See supplementary figure 2 for constructs design and SDS-PAGE of purified proteins. N1/09 hybrid means residues of loops 01 and 23 of N1/2009 grafted to mSN1 (H5N1/2021) scaffold. N1/19 hybrid means loops 01 and 23 of N1/2019 grafted to mSN1 (H5N1/2021) scaffold. The expression yield of NA proteins and their hybrid forms is included in the 2^nd^ column. N1/19 protein expression was undetectable to low in ExpiCHO cells in two instances. Size-exclusion chromatography graphs are shown in the 3^rd^ column. Elution volume of 10-14 mL indicates tetrameric form of the protein. MUNANA and ELLA activity of the NA proteins are in 4^th^ and 5^th^ columns with EC_50_ (effective concentration 50%) and AUC (area under curve) values, and the nanoDSF thermal melting temperature is in final column. The sharp narrow peak and the higher melting temperature indicate higher protein stability. **b) Epitope specificity has been transferred with loops.** Human monoclonal antibodies, previously published and some new, were titrated for ELISA binding of NA proteins. Areas under the curve were ranked after normalisation with one of the strongest binding mAb (see Supplementary Fig. 6a) ‘+++’ denotes >70% binding, ‘++’ 40-70% binding, ‘+’ 10-40%, and ‘-’ <10% as a non-binder. Loops cross-reactive mAbs AG7C, AF9C, Z2B3 and 1G01, all defined by crystal structures, show full binding to all NA proteins. Seven mAbs (NmAb) do not bind mSN1 but bind to N1/09, N1/19 and their hybrid forms. NmAb-03 is specific to N1/19 surface Loops. mAb CD6 is a scaffold dependent mAb that shows binding to N1/19 hybrid protein but does not bind the N1/19 protein on the cell surface (see Supplementary Fig. 5b). mAb Z2C2 is specific for mSN1 and N1/09, hence did not bind N1/19 or its hybrid form.

There was a substantial increase in protein yield for both N1/09 and N1/19 hybrid proteins (Fig. 3a). These hybrids expressed as tetramers with negligible aggregation, as demonstrated by size-exclusion chromatography (Fig. 3a) and a BS3 cross-linked tetrameric complex of ∼250 kDa resolved on SDS-PAGE (Supplementary Fig. 3b). The hybrids were enzymatically active in both small substrate MUNANA (580 Da) and large substrate fetuin (49 kDa) based assays. The N1/09 hybrid showed an 8°C improvement in its melting temperature T_m_ to 64.4°C compared to the N1/09 protein (56.4°C), while the T_m_ of the N1/19 hybrid (72.4°C) surpassed the H5N1 mSN1 scaffold donor (68.7°C). The 4.3°C difference in melting temperature between the mSN1 protein (68.7°C) and the N1/09 hybrid (64.4°C) may indicate that the sequence variation in the loops also contributes to stability of the tetramer.

Binding by mAb CD6 ^16,20^ was also informative as CD6 binds across two protomers of the tetrameric NA and makes contacts with both loop and scaffold residues (Supplementary Fig. 5a). Ellis et al have shown that binding of CD6 is dependent on the formation of compact closed conformation tetramers, whereas a more open conformation N1 2009 tetramer failed to bind CD6 ^16^.

Further, we compared the binding of CD6 to N1/09-VASP (with human vasodilator stimulated phosphoprotein tetramerisation domain) and N1/09-TB [with tetrabrachion (TB) tetramerisation domain] proteins expressed in ExpiCHO cells and found that CD6 recognised only the latter (Supplementary Fig. 5b). CD6 bound better to N1/09 infected cells than to the recombinant N1/09-TB protein (Fig. 3b, Supplementary Fig. 6a). This aligns with Ellis *et al*.’s findings that the 2009 N1 protein (with a VASP tetramerisation domain) predominantly formed open tetramers that failed to bind CD6. Our recombinant N1/09 protein with the TB tetramerisation domain may have formed a mixture of open and closed tetramers, also suggested by the appearance of aggregates on size-exclusion chromatography and the relatively low T_m_ of 56.4°C (Fig. 3a). In contrast, CD6 bound strongly to the N1/09 hybrid protein, which had a higher T_m_ of 64.4°C. CD6 also bound strongly to mSN1 scaffold donor.

The CD6 epitope was lost on seasonal H1N1 viruses isolated after 2015 (unpublished observation) so CD6 bound to cells infected with N1/09 but not N1/19 virus, but CD6 bound to the N1/19 hybrid protein strongly, suggesting that CD6’s binding to the N1/19 hybrid depended on the scaffold residues donated by the msN1 protein, and that the N1/19 hybrid protein had formed a compact tetramer compatible with the N1/19 hybrid’s higher T_m_ of 72.4°C (Figure 3a).

### N1/09 and N1/19 hybrid proteins retained epitope specificity and enzyme activity

Next, we assessed the NA hybrid proteins for the epitopes recognised by monoclonal antibodies isolated from humans infected with influenza virus or vaccinated with inactivated seasonal influenza vaccine. A comprehensive analysis was done using a collection of twenty-five human mAbs, isolated by us or synthesized in the laboratory based on published sequences (see Methods; Supplementary Fig. 3a). Results from the binding titration of fifteen mAbs on mSN1 and hybrid proteins and NA on the virus-infected cell membrane are depicted in Fig. 3b. These included a set of five broadly reactive mAbs, including four, 1G01 ^13^, Z2B3 ^10^, AG7C and AF9C (^21^ and personal communication with Dr. Yan Wu, Capital Medical University) with crystal structures showing binding to the L01 and L23 loops, and nine strain specific mAbs that distinguished between loop donor and recipient. Cross-reactive mAbs showed full binding to all proteins - mSN1, loop-donors, and hybrid proteins. Seven mAbs which did not bind the mSN1 protein but bound N1/09, also bound to the N1/09 hybrid protein, showing their specificity for the transferred loops. Similarly, NmAb-03, displaying specific binding to N1/19 NA on infected cells, exclusively bound to the N1/19 hybrid protein. By contrast, mAb Z2C2 which is specific for mSN1 and N1/09, bound neither N1/19-infected cells nor the N1/19 hybrid protein.

These results showed that for a wide collection of cross reactive and specific mAbs, isolated from influenza infected or vaccinated humans, binding was retained on the hybrid proteins. Together these results suggested that the tertiary structure of the loops had been retained after grafting onto the msN1 scaffold.

NA inhibiting drugs, oseltamivir and zanamivir, inhibited the NA activity of the hybrid proteins (Supplementary Fig. 3c). Similarly, mAb 1G01, a known catalytic site targeting mAb also inhibited the activity of mSN1 and N1/09 hybrid but not N1/19 hybrid. This result was expected since the mAb 1G01 lost its inhibitory activity on 2019 H1N1 seasonal influenza ^12^ due to the substitution N222K (N1 numbering) located within the loop B3L01 (unpublished observation, Fig. 3b and Supplementary Fig. 3c). While enzyme inhibition was lost, detectable binding of 1G01 was retained to N1/19 cells and N1/19 hybrid proteins (Fig. 3b, Supplementary Fig. 6a).

### mSN1, N1/09 hybrid and N1/19 hybrid crystal structures are nearly identical

To assess whether loop grafting influenced the local or overall structure of the neuraminidase, we determined high resolution crystal structures of mSN1, the N1/09 hybrid, and the N1/19 hybrid (Fig. 4, Supplementary Table 3). Overlay analysis revealed that N1/09 hybrid and N1/19 hybrid were nearly identical in structure to mSN1, with overall RMSD (root-mean-square deviations) of <0.25 Å across equivalent Cα atoms upon overlay (Fig. 4b) ^22^. This is consistent with previous findings, where N1 neuraminidases within a subtype typically exhibit an RMS deviation between 0.2−0.4 Å ^17,18^. All segments of the scaffold region in both NA hybrids superposed extremely well onto mSN1, with RMS deviation values of less than 0.5 Å between the most distantly aligned Cα residue pairs. Minor deviations were observed in the B1L23 (150-loop) and B6L23 (430-loop) regions bordering the active-site cavity, with the RMS deviation distances of up to ∼1.0 Å between aligned Cα residue pairs. The remaining loops showed RMS deviation values of less than 0.5 Å.

**Fig 4:**
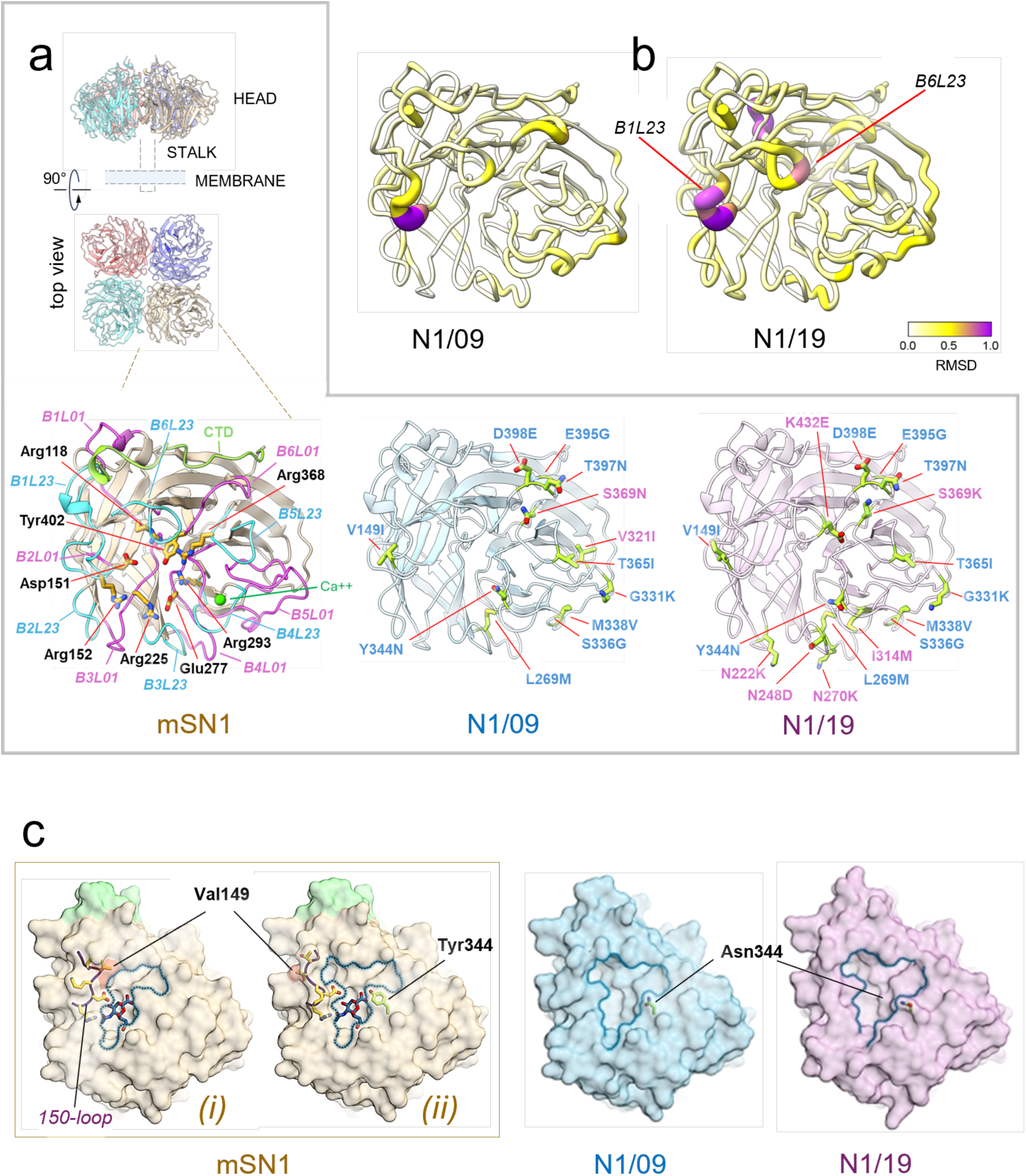
The structure of the influenza virus neuraminidase is strictly conserved in the loops-grafted hybrid proteins. **a) X-ray crystal structures.** *(top)* The influenza virus neuraminidase is shown in the native tetrameric form, an assembly observed in all reported crystal structures. A schematic representation of the stalk and a membrane is shown. *(bottom)* Structures of each crystallized constructs viewed from above the β-propeller fold (cartoon representation). *(left)* Crystal structure of a protomer of the mSN1 neuraminidase. The β-strands of the H5N1/21 protomer (mSN1, left) are colored pale brown, with the B6L01 and B6L23 loops colored magenta and cyan blue, respectively. The C-terminal domain (CTD) is shown in pale green, and the calcium ion is depicted as a green sphere. Eight highly conserved residues in the catalytic site (R118, D151, R152, R225, E277, R293, R368, and Y402, N1 numbering based on msN1) are shown as yellow sticks. *(middle)* Crystal structure of a protomer of the N1/09 Loops-mS hybrid (pale blue). *(right)* Crystal structure of a protomer of the N1/19 Loops-mS hybrid (pale violet). B6L01 and B6L23 residues grafted into the N1/09 and N1/19 hybrid proteins are shown as yellow-green sticks. Residues identical in the N1/09 and N1/19 hybrids are labelled in blue; residues differing between these hybrids are labelled in violet. **b) Mapping structural differences between N1/09 and N1/19 hybrid structures.** Local root-mean square (RMS) deviations between equivalent Cα pairs are mapped following the overlay of N1/09 *(left)* and N1/19 *(right)* onto the crystal structure of msN1. The b-propeller of msN1 is shown in a putty tube representation, with color and radius reflecting the local RMS deviation values between equivalent Cα pairs. RMS deviations are generally below 0.5 Å, with modestly higher values in the inherently flexible B1L23 and B6L23 loops. **c) Active site cavities.** The msN1 surface is colored brown, except for the C-terminal loop region (green). The N1/09 and N1/19 surfaces are in blue and violet, respectively. In mSN1, the rim of the cavity containing the active site is traced with a blue dashed line. The position of the active site is denoted by a sialic acid molecule (grey-blue sticks), taken from a superposed, related structure (PDB ID: 2BAT). The msN1 structure revealed differences among the protomers of the tetramer at the active site entrance. Within the same tetramer, protomers with a relatively narrow cavity (*i*) combine with protomers showing a wider entrance (*ii*). This difference is dictated by the trajectory of the B1L23 loop (*150-loop*; dark blue with yellow side chains). The variation in trajectory between (*i*) and (*ii*) is most pronounced at Val149 (red surface). In contrast, the N1/09 and N1/19 hybrids, no noticeable differences exist between the protomer cavities, all of which closely resemble the wider msN1(*ii*) conformation, apart from minor, local widening of the rim induced by the substitution of Y344 with an asparagine residue (shown as light-green sticks).

While the scaffold is structurally conserved across the three resolved structures, each possess unique active site conformations, which are primarily influenced by the L01 and L23 loops surrounding the active site cavity. Analysis of mSN1, N1/09 hybrid, and N1/19 hybrid structures revealed some variability in active site conformations, with the greatest variability existing between the cavities of two msN1 molecules within the asymmetric unit of the crystal, reflecting different trajectories of the B1L23 (150-loop) (Fig. 4c). Additionally, we observed that the Y344N substitution in the N1/09 and N1/19 hybrids slightly widened the enzymatic cavity compared to msN1.

The structural and monoclonal antibody binding data thus showed that the twelve or sixteen substitutions in loops matching the 2009 and 2019 seasonal N1s grafted onto the avian N1 scaffold did not affect the protein fold. The NA hybrid proteins exhibited a high degree of structural conservation, consistent with all previously published crystal structures of N1 neuraminidase proteins ^17,23^.

### N1/09 and N1/19 hybrid proteins are immunogenic and elicit NA enzyme inhibiting (NAI) antibody responses

We have previously shown that recombinant NA presented on a mi3 particle is strongly immunogenic at doses as low as 0.1 μg ^24^, in contrast to 10-100 fold higher doses of free NA protein often described in the literature ^4^. Our recombinant NA proteins were coupled onto mi3 virus-like particles (NA-VLP) via the covalent bond between the SpyTag on the NA protein and SpyCatcher003 sequence on the mi3 protein (Supplementary Fig. 3d). Mice were immunised with 0.5 μg NA-VLP adjuvanted with 1:1 vol/vol AddaVax^TM^ (squalene-based oil-in-water nano-emulsion). Intramuscular immunisations were administered twice at three-week intervals, and sera were harvested three weeks post-booster dose to assess the antibody response. Neuraminidase activity inhibition (NAI) IC_50_ (inhibitory concentration 50%) titres were measured using a fetuin-based enzyme-linked lectin assay (ELLA).

All NA proteins were immunogenic in mice, generating binding and NA inhibiting antibodies (NAI) to themselves (Fig. 5a). mSN1 elicited NAI antibody titres against itself that cross inhibited viral N1/09 NA. However, N1/09 protein did not induce a cross-reactive response against mSN1, although the N1/09 hybrid did elicit antibodies that inhibited mSN1. This difference may have been due to improved structural integrity of the N1/09 hybrid, or possibly due to induction of inhibitory antibodies to the mSN1 scaffold.

**Fig 5:**
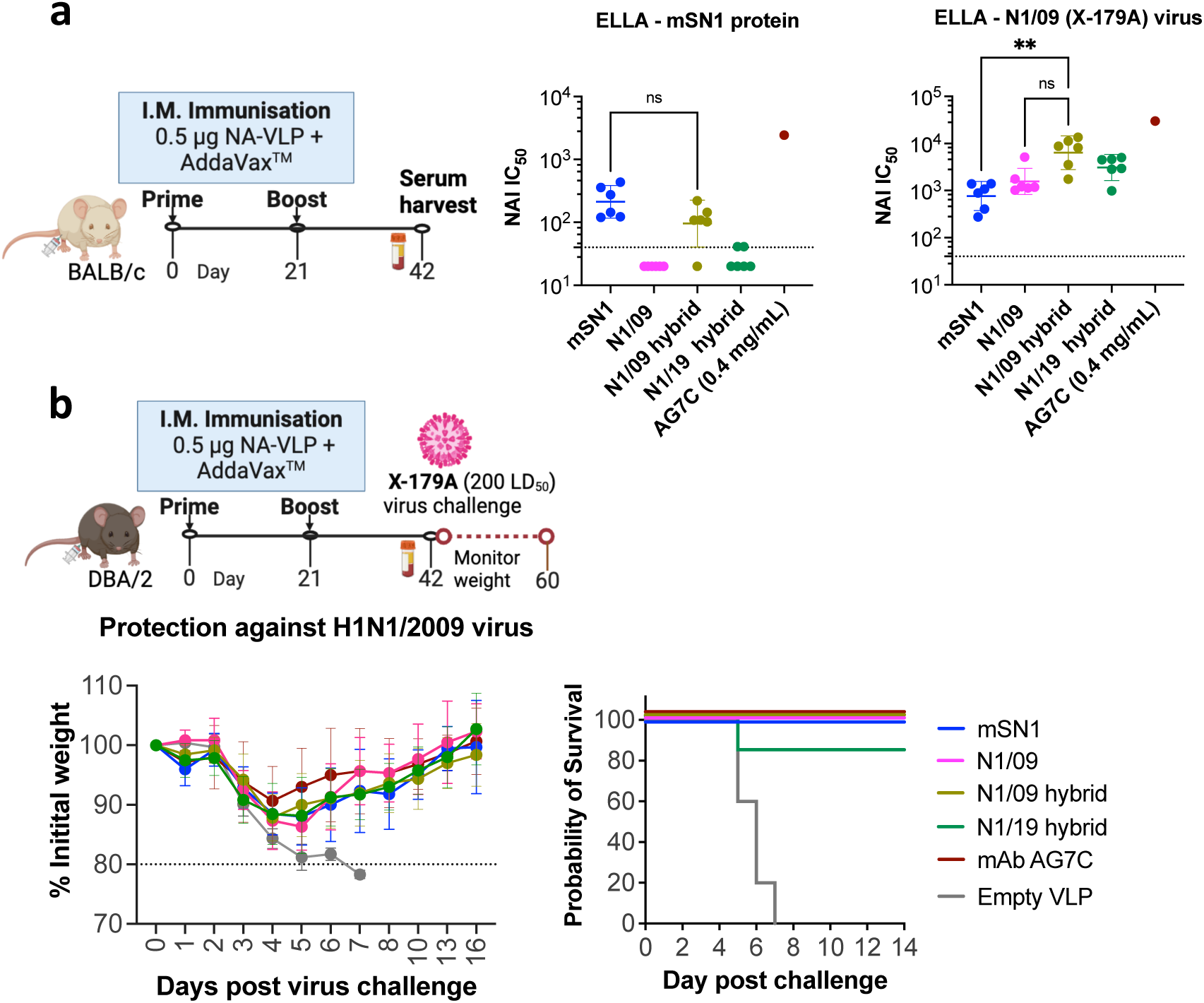
NA-hybrid proteins are immunogenic and provide *in vivo* protection against virus challenge. **a) Immunogenicity of NA hybrid proteins.** BALB/c mice (n=6/group) were immunised with 0.5 μg NA coupled to the mi3 virus-like particles (NA-VLP) adjuvanted with 1:1 vol/vol AddaVax^TM^ (squalene-based oil-in-water nano-emulsion). Intramuscular immunisations were done twice at the interval of three weeks and sera were harvested three weeks post booster dose to assess the antibody response. Neuraminidase activity inhibition (NAI) IC_50_ titres measured using fetuin-based Enzyme-Linked Lectin Assay (ELLA) are shown as a separate dot for each mouse. mAb AG7C was used as a positive control. Pooled sera to unconjugated mi3 VLP was negative control and showed no inhibition at 1:40 dilution (not included here). Geometric mean with 95% confidence interval is shown. **b) Protection against virus challenge.** DBA/2 mice (n=6/group) were immunised as above and were challenged with intranasally administered 200 LD_50_ of H1N1/2009 (X-179A) virus. Weights were monitored for two weeks. Loss of ≥20% initial weight was considered an endpoint. mAb AG7C (10 mg/Kg prophylaxis) was used as a positive control. Empty VLP pooled sera was a negative control and mice reached the endpoint within day 5-7 post virus infection. The ELLA NA inhibition graphs of these pooled sera are shown in Supplementary Fig. 7. Figures were made using GraphPad Prism v10. Kruskal-Wallis test was used for statistical analysis. ns: non-significant (p-value>0.05), ** means p-value<0.005. Kaplan-Meier survival analysis was done with Logrank Mantel-Cox test for comparison.

The mSN1 and the N1/19 hybrid both induced strong inhibitory antibodies to themselves, but the sera to mSN1 failed to cross-inhibit the enzyme activity of N1/19-like virus (Supplementary Fig. 7b,c) and the N1/19 hybrid protein did not generate a cross-inhibitory NAI antibody response to mSN1. So, in this case the common mSN1 scaffold did not generate cross-inhibitory antibodies. N1/19 had four significant additional amino acid substitutions (see discussion) compared to N1/09 in the transferred L01 and L23 loops which may have been responsible for the loss of cross reactivity between 2019 N1 with the 2021 avian mSN1. However, the N1/19 hybrid did generate antibodies that cross-inhibited N1/09 virus, so all four immunogens mSN1, N1/09, N1/09 hybrid and, N1/19 hybrid induced antibodies that could inhibit H1N1/2009 neuraminidase.

### Murine Challenge Studies

We challenged immunised DBA/2 mice with a lethal dose of H1N1/2009 (X-179A) virus and monitored weights for at least 14 days. Mice immunised with empty VLP lost ≥20% initial weight and hence were culled within 5-7 days post-infection. Vaccination with all the tested NA proteins mSN1, N1/09, N1/09 hybrid and N1/19 hybrid protected mice from severe weight loss on challenge with X-179A. These results mirrored several studies in the literature which showed that immunisation with the 2009 N1 could provide at least partial protection in mice and ferrets to the avian H5N1 challenge ^25–27^, and is compatible with the well characterised human mAbs that cross-inhibit broadly within the N1 subtype (^21^, reviewed in ^5,6^).

In this case all four immunogens induced inhibitory antibody to 2009 viral NA, resulting in protection from challenge with H1N1/2009 virus.

Although there is broad cross reactivity of antisera within the N1 subtype ^25–27^, reviewed in ^4^, it is not universal ^14,28^ as demonstrated above with mSN1 and the N1/19 hybrid (Supplementary Fig. 7b,c) which differed by 16 amino acids in the transferred loops, compared to 12 differences for the N1/09 hybrid (Figure 2). We noted that the Cambridge strain of H1N1 A/PR/8/1934 and mSN1 differed by 18 residues in the L01 and L23 loops which may be sufficient to prevent cross-inhibition. We then exchanged the L01 and L23 loops between A/PR/8/34 and mSN1 to look for correlation between antibody cross-reactivity and protection with the source of the L01 and L23 loops.

### Loop transfer between two distant N1 NAs: H5N1 A/mute swan/England/053054/2021 (mS) and H1N1 A/PR/8/1934 Cambridge Strain (PR8)

mSN1 showed sufficient cross-reactivity to N1/09 to protect mice against virus challenge. Therefore, we performed loop transfer between mSN1 and PR8N1, which differ by 18 residues within the L01 and L23 loops and show no or minimal cross-reactivity, to assess the loop-specific protection. We exchanged the L01 and L23 top surface loops between mS and PR8 that differ by 18 residues (Fig. 6a, Supplementary Fig. 8). Following the nomenclature above we will refer to these as the PR8 hybrid (PR8 Loops-mS) comprised of Loops 01 and 23 from PR8 combined with the mSN1 scaffold and mS hybrid (mS Loops-PR8) comprised of loops 01 and 23 from mS combined with the PR8 scaffold. Unlike the N1/09 and N1/19 proteins, wild-type PR8 N1 expressed well (∼105 mg/L; T_m_ 55.8°C) as tetramers with minimal aggregation (Refer to SDS-PAGE in Supplementary Fig. 3b). The PR8 hybrid expressed at a nearly equal yield (108 mg/L; T_m_ 58.1°C) and showed minimal aggregation (Fig. 6b). Conversely, the mS hybrid gave a lower yield at (∼22 mg/L; T_m_ 64.3°C) but also showed a single peak in melting temperature and assembled into a tetramer with minimal aggregation (Fig. 6b).

**Fig 6:**
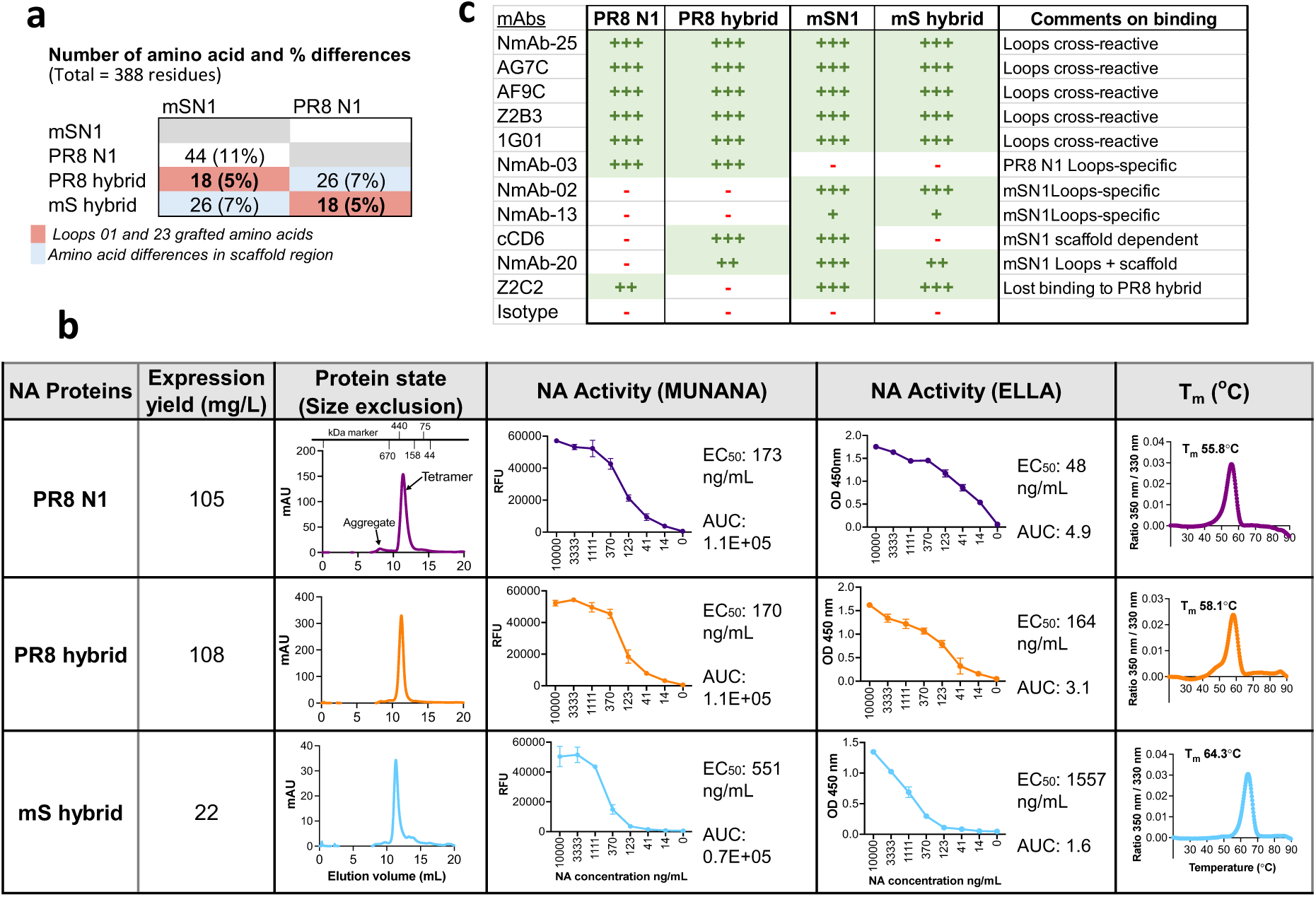
Loop grafting between two distant N1 NAs: H5N1 A/mute swan/England/053054/2021 (mS) and H1N1 A/PR/8/1934 (PR8) **a)** Number of amino acid differences between mSN1 and PR8 N1 and their loop grafted hybrids are shown. Eighteen dissimilar residues (5%) within Loops 01 and 23 were grafted to make the hybrid proteins. **b) Characteristics of proteins.** NA proteins were expressed in a transient mammalian ExpiCHO expression system. The expression yield of NA proteins and their hybrid forms is included in the 2^nd^ column. Size-exclusion chromatography graphs are in the 3^rd^ column. Elution volume of 10-14 mL indicates tetrameric form of the protein and 14-15 ml indicates trimeric or dimeric nature of the protein. MUNANA and ELLA activity of the NA proteins are in 4^th^ column and the nanoDSF thermal melting temperature is in the final column. The sharp narrow peak and the higher melting temperature indicate the higher protein stability. **c) Epitope specificity has been transferred with Loops with a few exceptions.** Antibodies, previously published and some new, were titrated for ELISA binding of NA proteins. Area under curve was ranked after normalisation with one of the strongest binding mAb (Refer to Supplementary Fig. 6b for binding titration data).) ‘+++’ denotes >70% binding, ‘++’ 40-70% binding, ‘+’ 10-40%, and ‘-’ <10% as a non-binder. Loops cross-reactive mAbs recognised both mSN1 and PR8N1 and their hybrid proteins. PR8N1 Loops specificity is shown by NmAb-03. mAb CD6 is a major scaffold dependent mAb and showed binding to PR8Loops-mS but not the PR8 and mSLoops-PR8 (see Supplementary Fig. 6b). Similarly, NmAb-20 is a mSN1 Loops + scaffold dependent mAb. mSN1 Loops specificity is shown by NmAb-02 and NmAb-13.

**Fig 7:**
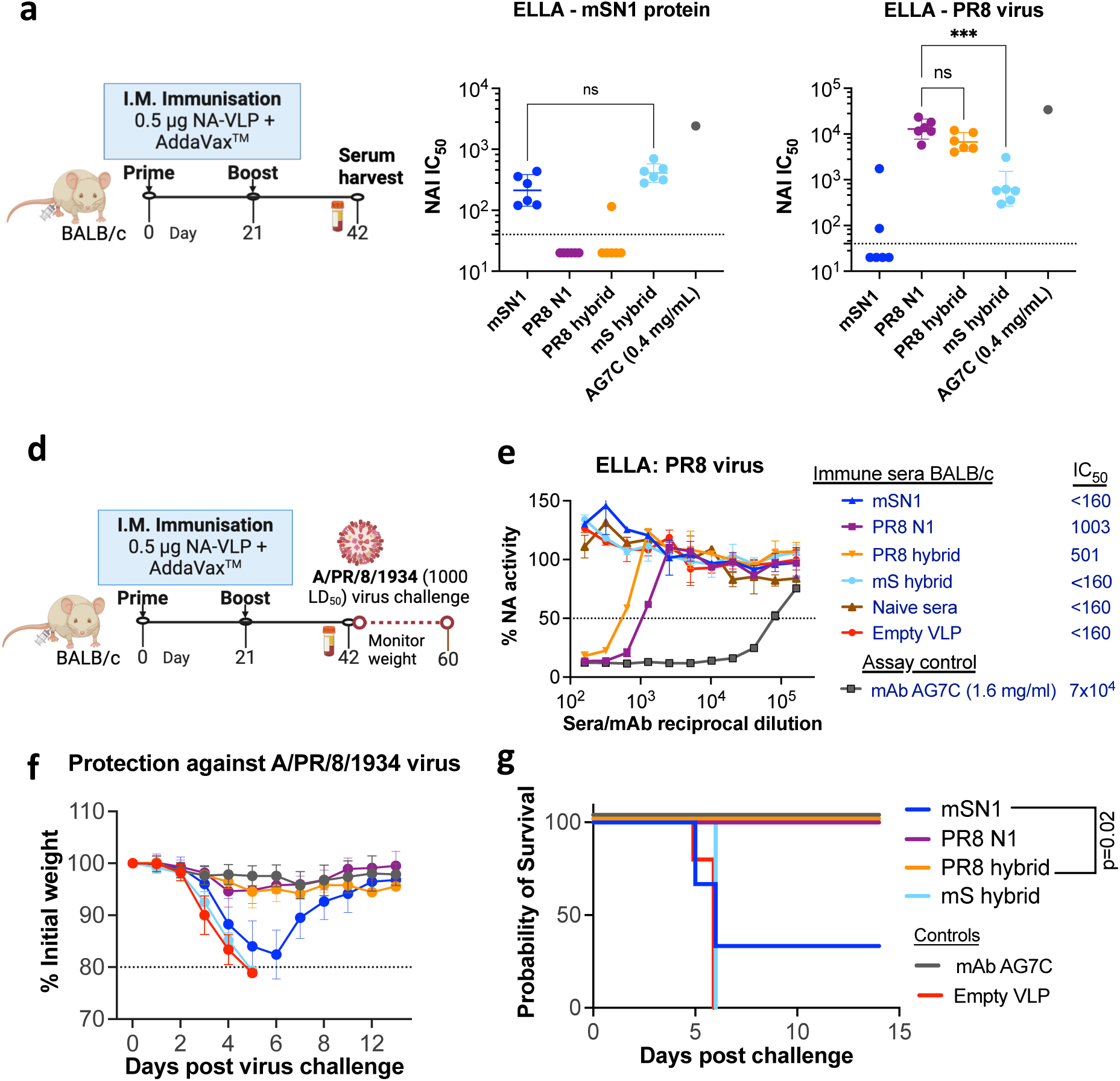
NA hybrid proteins elicited loop-specific NA inhibiting antibodies and provided loop-specific protection *in vivo* against virus challenge. **a,b,c) Immunogenicity of NA hybrid proteins.** BALB/c mice (n=6/group) were immunised with 0.5 μg NA coupled on to the mi3 virus-like particles (NA-VLP) adjuvanted with 1:1 vol/vol AddaVax^TM^ (squalene-based oil-in-water nano-emulsion). Intramuscular immunisations were done twice at the interval of three weeks and sera were harvested three weeks post booster dose to assess the antibody response. Neuraminidase activity inhibition (NAI) IC_50_ titres measured using fetuin-based Enzyme-Linked Lectin Assay (ELLA) are shown as a separate dot for each mouse. mAb AG7C was used as a positive control. Empty VLP pool sera were negative controls and showed no inhibition at 1:40 dilution (not included here). Geometric mean with 95% confidence interval is shown. **d,e,f,g) Protection against virus challenge.** BALB/c mice (n=6/group) were immunised as above. Pooled sera antibodies were assessed in ELLA assay before virus challenge (e) and IC_50_ values shown are sera reciprocal dilution. Mice were challenged with intranasally administered 1000 LD_50_ of PR8 virus (Cambridge strain, 10^4^ TCID_50_). Weights were monitored for two weeks. Loss of ≥20% initial weight was considered an endpoint. mAb AG7C (10 mg/Kg prophylaxis) was used as a positive control. Empty VLP pool sera were negative controls and mice reached the endpoint by day 6 post virus infection. Importantly, immunogens with PR8 Loops protected 100% mice from virus challenge and mS Loops did not. Mean and standard deviations are shown. Figures were made using GraphPad Prism v10. Kruskal-Wallis test was used for statistical analysis. ns: non-significant (p-value>0.05), *** denotes p-value<0.0005. Kaplan-Meier survival analysis was used with Logrank Mantel-Cox test for comparison.

### Epitope specificity between loop-exchanged mS and PR8 hybrid proteins

We assessed the epitope specificity of these proteins, following the methods applied for N1/09 and N1/19 hybrids (Supplementary Fig. 6b). We titrated 17 mAbs for binding to the wild-type and hybrid NA proteins, with results for representative 11 mAbs shown in Fig. 6c. Ten cross-reactive mAbs showed full binding to all proteins - PR8 N1 and mSN1 and their loops-exchanged hybrids. Among specific mAbs, NmAb-03 specifically bound to PR8 N1 and the PR8 hybrid but did not bind to mSN1 or the mS hybrid, thus showing specificity for the PR8 loops. Similarly, NmAb-02 and -13 bound to mSN1 and the mS hybrid but not to PR8 N1 or the PR8 hybrid, thus showing specificity for the mS loops. CD6 is a scaffold-dependent mSN1 binder and did not bind to PR8 N1 NA. CD6 maintained its binding to mSN1 scaffold and to the PR8 hybrid but not to the mS hybrid, indicating its dependence on scaffold residues from the H5N1 mS donor (Supplementary Fig. 5a).

mAbs NmAb-20 and Z2C2 were exceptions. NmAb-20 bound mSN1 and the mS hybrid suggesting that it was mSN1 loops specific. However, in addition it *gained* binding to PR8 hybrid. mAb Z2C2 bound to both PR8 and mSN1, but lost binding to PR8 hybrid, implying that this epitope had been lost in the PR8 hybrid protein. These two exceptions out of 17 mAbs studied in detail suggest that the transferred loops may show a limited amount of antigenic difference to either donor, perhaps at the margins of the loops.

### mS and PR8 hybrid proteins elicited loop-specific NA inhibiting (NAI) antibodies and provided loop-specific protection *in vivo* against virus challenge

BALB/c mice were immunised with 0.5 μg NA-VLP adjuvanted with AddaVax^TM^ (Fig. 6a), as previously described for N1/09 in vivo experiments. mSN1 and mS hybrid proteins generated equivalent NAI sera titres towards mSN1, whereas PR8 N1 and the PR8 hybrid elicited no detectable titres to mSN1 (Fig. 6b). Similarly, PR8 and PR8 hybrid generated equivalent titres against PR8 virus (Fig. 6c). Interestingly, the mS hybrid elicited a low NAI titre (albeit ∼10-fold lower compared to PR8 N1; p=0.0009) against PR8 virus, implying that in this combination some inhibitory antibody may have been generated either to the PR8 scaffold or to conserved epitopes in the L01 and L23 loops.

In an independent experiment, we challenged immunised mice with a lethal dose of PR8 virus (Cambridge strain) (Fig. 6d). Weight-survival curves showed that 6/6 mice immunised with PR8 NA and the PR8 hybrid NA survived without weight loss, whereas all mice vaccinated with the mS hybrid reached the endpoint, similar to the negative control group immunised with empty VLP (Fig. 6f,g). For mice immunised with mSN1 wild-type NA, 2/6 mice survived but with significant weight loss. These experiments demonstrated that survival matched pre-exposure to vaccine presenting the PR8 L01 and L23 loops. The serology of vaccinated mice showed the same pattern (Fig. 6e). PR8 and the PR8 hybrid vaccines generated inhibitory titres to PR8 virus NA, but mSN1 and the mS hybrid elicited no detectable NAI titres to PR8 virus (<160 in this experiment). The majority of the NAI serum titres were generated against the loops and this matched protection in this pair of NAs that differed by 18 residues in the L01 and L23 loops.

## Discussion

Our Loop-grafting concept is straightforward and may be broadly applicable within other NA subtypes. The concept is based on three simple assumptions: 1) the majority of protective epitopes for the antibody response are present in the L01 and L23 loops on the top surface of the NA head as defined by Varghese and Colman ^7,29^, 2) the stability of the NA tetramer depends mainly on interactions between monomers encoded in the remainder of the NA sequence (the “scaffold”), and 3) the structure of the loops will be maintained when grafted from one scaffold to another, at least within an NA subtype ^30–32^.

### The role of the tetramerisation domain

In preliminary experiments we compared the VASP and Tetrabrachion (TB) tetramerisation domains for expression and immunisation with the 2009 N1 protein. We found that the VASP domain supported the formation of a NA tetramer but the tetrameric protein did not bind the CD6 antibody that binds across two monomers (Supplementary Figure 5b), confirming results of Ellis *et al*. ^16^. By contrast the N1/09-TB did bind CD6 and protected mice against matched viral challenge, as shown here. This result suggested that, while the VASP-linked 2009 N1 does form tetramers as defined by size-exclusion and cross-linking, the protein may not be in an optimal state for vaccination. We therefore have used the tetrabrachion domain to form tetramers throughout this report.

### The basis for stable tetramer formation

The hybrid proteins we produced all appeared to form stable tetramers as defined by SEC, BS3 crosslinking and melting temperature. They were all active in ELLA (large substrate) and MUNANA (small substrate) enzyme activity assays and were inhibited by standard small molecule inhibitors. We tested a wide range of virus specific and cross-reactive human mAbs and with few exceptions the antibodies bound to the hybrid proteins. In addition, the CD6 antibody, that binds across two monomers and only to fully formed tetramers ^20^, bound to our hybrid proteins held together by the TB tetramerization domain. Finally, the crystal structures of our hybrid N1 proteins confirmed that the grafted loops had retained their expected conformations. Together these data suggested that the hybrid proteins had folded correctly and, in most pairs, combined the expression and stability characteristics of the scaffold donor with the antigenic properties of the loop donor.

Ellis *et al*. defined forty-four residues that form contacts between N1 2009 monomers that could be involved in maintaining the stability of a tetramer ^16^. In their final design of the stabilised sNAp-155 N1 2009 NA they replaced ten of these forty-four amino acids, to obtain a stable 2009 N1 tetramer supported by a VASP tetramerization domain (Supplementary Fig. 9). In our hybrids, tetramerized with TB, these ten residues are completely conserved between the scaffold and loop donors, so the source of stability for our hybrid proteins must lie elsewhere.

### The distribution of epitopes on Neuraminidase

Varghese, Laver, and Colman described the structure of L01 and L23 loops on the top surface of N2 NA and showed that these loops contribute largely to the active site, notably including seven of eight conserved residues in the catalytic site and eight of eleven conserved framework residues that support the site ^7,8^. Also, in the L01 and L23 loops surrounding the active site they noted multiple variable residues that vary seasonally associated with antigenic drift, and that can be selected for viral resistance by monoclonal antibodies *in vitro*. Resistance mutations selected for by three N2 murine mAbs ^33^ and N9 mAbs were within loops B3L01, B5L01 and B5L23 ^8^. Over the last forty years many similar studies have been done with murine and human monoclonal antibodies to various NAs. In supplementary figure 1 we have collected a set that identifies thirty-one sites of amino acid substitution selected by monoclonal antibodies in independent experiments. Twenty-five of these are in Loops 01 and 23, with three additional sites in immediate neighbor positions to these loops (90% altogether). Three additional sites of selection can be assigned to the underside of the NA head at the interface of stalk and head (position 88), B4 Loop 12 (position 285), B4 Loop34 (position 309). In addition, a recent study of human sera identified position 386 on B5 loop 34 as responsible for an antigenic change between N1/1977 and N1/1986 ^34^.

Crystal structures of bound Fab fragments give further information on the footprints of protective antibodies (Supplementary Fig. 2). The majority describe binding to the L01 and L23 loops, with recent examples also confirming binding to the side (CD6, NA-22) and underside of NA ^20,35–38^. It is important to note that loops 01 and 23 include a part of epitopes that have been described in the literature as side, lateral, or underside (see mAbs NDS.1, NDS.3 and CD6 in Supplementary Fig. 2). The majority of antibodies defined by crystallography bind within the surface of a single monomer, with recent structural evidence that a few mAbs binds across two monomers ^20,37^. Finally, several structures of antibodies that bind within the active sites of a broad range of NAs have been described, all of which contact the L01 and L23 loops ^10–13^. From these data we suggest that most antibodies generated by NA that are likely to be protective bind to the L01 and L23 loops, while a minority bind to epitopes on the underneath and side of the NA head that are not included within the L01 and L23 loops.

### Definition of Loops

To align the top surface loops in N1 proteins, we relied on loop annotations from the N2 NA structure resolved to 2.9 Å by Varghese *et al.* as a reference ^7^. The loop definitions in the structure later resolved to 2.2 Å differed slightly ^29^ Supplementary Fig. 4). We suggest that loop annotations for a grafting experiment should not be rigid and requires some judgement, particularly for residues at the margins of loops, taking into consideration solvent accessibility in crystal structures, evidence for evolutionary selection, viral resistance mutations selected with mAbs *in vitro* (Supplementary Fig. 1), and defined crystal structures of bound antibodies (Supplementary Fig. 2). For instance, in our N1/19 hybrid design, residue N200S, could have been considered as part of loop B2L23 (^29^ and Fig. 2a). In addition the C-terminal domain (CTD) contributes to surface residues and may be suitable for grafting, since residues within this section of NA have been selected by mAbs *in vitro* and clearly evolve over time ^8^.(Supplementary Fig. 1,2). However, the CTD of N1 was conserved in all of our examples.

### Vaccination with hybrid NAs

Our NA vaccination strategy is based on our earlier evidence that linking tetrameric NAs to the mi3 vaccine-like particle (valency of sixty) via a SpyTag/SpyCatcher covalent linkage results in enhanced immunogenicity and dose sparing, with doses as low as 0.1 μg of NA protein able to induce NAI antibody ^24^. This compares well to the higher doses of pure protein used in the majority of studies of NA immunity (reviewed in ^6^). In the present experiments we opted for two doses of 0.5 μg of mi3 linked NA that gave full protection in preliminary experiments (not shown), comparable to recent results from Pascha *et al.* ^39^.

In our first set of grafting experiments between N1/2009 and mSN1 (H5N1 2021) we found that while protein expression was greatly improved, antisera from animals were cross reactive for NA inhibition between the scaffold H5N1 donor, the hybrid (with 12 amino acid replacements in the L01 and L23 loops) and the 2009 Loop donor. In addition, all immunised animals were protected from challenge with 2009 H1N1 virus. These results were consistent with evidence that within the N1 subtype there is broad cross-reactivity for antibodies and that immunisation with 2009 N1 provides at least partial protection against an avian H5N1 (reviewed in ^6^). Cross-reactivity could have been for epitopes anywhere within the NA structure.

The sequence difference in the loops between 2019 N1 donor and the H5N1 recipient was greater (16 amino acids), sufficient to provide an antigenic distance that prevented cross inhibition by antisera from animals immunised with these two NAs. The NA inhibitory activity of the antisera was predominantly specific for the L01 and L23 loops. N1/19 had four significant additional amino acid substitutions compared to N1/09 in the transferred L01 and L23 loops: N222K (B3L01), N244D (B3L23), N270K (B4L01) and K432E (B6L23). All four of these residues have been selected for resistance by inhibitory mAbs *in vitro* (see Supplementary Fig. 1), or form contacts with mAbs demonstrated in crystal structures (Supplementary Fig. 2), so these residues are likely to have been responsible for the loss of cross reactivity between N1/2019 and mSN1. We do not have a challenge model for the 2019 H1N1 virus, so we could not look for in vivo protection specific to the 2019 N1 loops.

The lack of cross inhibition between sera raised to the avian mSN1 and the seasonal N1/19 showed that cross reactivity of antisera to the N1 NAs is not universal, and reflects the results of Lu *et al.* ^14^ who found minimal cross reactivity of rabbit sera raised to a seasonal N1 from A/Beijing/262/1995 and an avian N1 from A/Hong Kong/ 483/1997 that we note differed by seventeen amino acids in the L01 and L23 Loops.

We found that the NA of mouse-virulent H1N1 virus A/PR/8/1934 (Cambridge strain) differed by 18 amino acids in the L01 and L23 loops from the mS H5N1 scaffold donor. We therefore prepared two further hybrids in which the loops were exchanged between these two NAs. The relevant hybrids induced NA inhibitory sera that were largely loop specific, and protection against a high dose challenge with A/PR/8/34 was now dependent on matched L01 and L23 loops. In this case antibodies to the scaffold could not have been protective, and protection correlated with the NA inhibition activity induced, mirroring the evidence from human challenge and vaccination studies (reviewed in ^4,6^).

Taken together these results suggest caution should be exercised in designing vaccination strategies that rely on cross-protection between seasonal NAs and future pandemic NAs.

### Limitations of the study

The three assumptions listed at the start of the discussion may not always hold. Clear examples exist of protective monoclonal antibodies that bind to epitopes in the NA scaffold, including those that bind epitopes on the side ^20,35,37^ and underneath of the head domain ^36^. If these formed the majority of the response to vaccination our approach would fail due to the first assumption. However, we have found that at least for the N1 subtype the transferred loops induce both NA inhibiting antibody and provide protection against viral challenge in the mouse. In addition, loops on the underside of NA could also be exchanged. The second assumption may also be sensitive to exceptions. Ellis *et al.* identified forty-four residues that may contribute to the contacts between NA monomers, ten of which we note are in loops L01 and L23 ^16^. If the latter were variable and turned out to be critical for stability of the tetramer our approach would fail. We note that one of the hybrids we produced (mS hybrid with PR8 scaffold) expressed at a lower level than either parent. Finally, the assumption that the tertiary structure of the loops is maintained after grafting may not completely hold as we have found occasional mAbs that alter their binding to the hybrid protein. However, the great majority of NA inhibitory and protective antibodies isolated from humans bound the four hybrid NAs we have made. In addition, crystal structures revealed that the transferred loops of 2009 and 2019 NA donors were virtually superposable on those of the H5N1 recipient.

While we acknowledge these potential shortcomings of our approach, we recommend the loop-grafting strategy as a simple and pragmatic method to improve the yield and stability of NA protein vaccine antigens. NA loop grafting can also serve as a useful starting point for analytical methods aimed at further improvement ^16,40^. Computationally-optimised sequences ^40,41^ designed for cross-protection might also be improved by focusing attention on the variation within the L01 and L23 loops.

### Impact

Purified NA has been safely administered to humans as a vaccine candidate ^1^ and NAI antibodies to NA have been shown to be an independent correlate of protection against influenza infection (reviewed in ^4^). Enhancing current vaccines with NA proteins could substantially boost their effectiveness. A major challenge has been the production of NA at optimal yields. In this study, we developed an innovative solution via loop-grafting to produce stable and immunogenic NA protein at high yields that addresses this critical issue. Moreover, this study presents an epitope-based vaccine design and protein engineering for generating protective immunity, which could potentially be applied to other antigens.

## Methods

### Cell lines and viruses

ExpiCHO (Chinese hamster ovary) cells (Thermo Fisher) were used for the production of NA proteins and monoclonal antibodies. They were handled according to the manufacturer’s protocol. MDCK-SIAT1 (Madin-Darby Canine Kidney cells stably transfected with human α 2,6-sialyltransferase, SIAT1) cells were used for the production of viruses, and for virus infection for epitope specificity assays^42^. Cells were maintained in D10 medium [Dulbeccoʹs Modified Eagle Medium (DMEM) supplemented with 10% (v/v) foetal calf serum (Sigma-Aldrich, F9665), 2 mM glutamine, 100 U/mL penicillin and 100 µg/mL streptomycin], with incubation in a humidified 5% CO_2_ 37 °C incubator. DMEM supplemented with 2 mM glutamine, 10 mM HEPES (N-2-hydroxyethylpiperazine-N’-2-ethanesulfonic acid), 0.1% BSA (Bovine serum albumin), 100 U/mL penicillin and 100 µg/mL streptomycin was used for virus growth and virus dilution. The medium is referred to as virus growth medium (VGM). All cell lines were tested to be mycoplasma-free.

Viruses were obtained from the Worldwide Influenza Centre, Crick Institute, UK and the National Institute for Biological Standards and Control, UK. Viruses were propagated by infecting a monolayer of MDCK-SIAT1 with ∼0.1 MOI (multiplicity of infection) virus for 1 h before replacing with VGM containing 0.5-1 µg/mL TPCK (L-1-tosylamido-2-phenylethyl chloromethyl ketone)-treated trypsin. Virus supernatant was harvested after 48 h.

### Monoclonal antibodies

Antibodies AG7C, AF9C, Z2C2, and Z2B3 were previously published by our laboratory ^21^. mAb 1G01 was produced in the lab using the antibody sequence obtained from PDB 6Q23 ^13^. Genes were synthesized by GeneArt (Life Technologies), cloned into antibody expressing vectors, and expressed in ExpiCHO cells. Similarly, mAb CD6 was synthesized using the sequence obtained from PDB 4QNP ^20^. The mAb is chimeric since the mouse variable region is cloned with the human constant regions.

NmAbs and 24-1C are anti-NA humans mAbs provided by our collaborator Prof. Kuan Ying Huang (National University Taiwan), which we produced in ExpiCHO expression system using their expression plasmids (see Supplementary Fig. 6). The study protocol and informed consent of human monoclonal antibody isolation were approved by the ethics committee at the National Taiwan University Hospital and Chang Gung Memorial Hospital and were carried out in accordance with the Declaration of Helsinki and Good Clinical Practice guidelines. Written informed consent was received from each adult prior to inclusion in the study.

### Neuraminidase protein expression and purification

Gene constructs (except for the N1/09 protein) were designed to have H7 HA signal sequence (MNTQILVFALIAIIPTNADKI), Strep-tag II (SAWSHPQFEK), G3S linker, 6xHis, S3GSG linker, SpyTag (AHIVMVDAYKPTK), G2SG4S linker, and tetrabrachion tetramerisation domain from *Staphylothermus marinus* ^19^, GGSGTG linker, and finally an NA ectodomain (82-469) at the C-terminus (Supplementary Fig. 3a). Sequences were synthesized as human-codon-optimized cDNAs by GeneArt (Life Technologies) and cloned into pcDNA3.1/-plasmid for transfection. Proteins were expressed in ExpiCHO expression system (Thermo Fisher). Briefly, ExpiCHO cells were cultured in a humidified Multitron Cell incubator (Infors HT) at 37 °C with 8% (v/v) CO_2_, rotating at 125 rpm, for at least 2 passages before transient transfection. The Max-titre protocol from the manufacturer was used and proteins were harvested on day 8/9 post-transfection. Culture supernatants were clarified by centrifugation at 3000 *g* for 10 min at room temperature, followed by filtration through a 0.22 µm filter (RapidFlow Nalgene). Proteins were purified from cell culture supernatants using Ni-NTA-sepharose (prepacked HisTrap HP column from Cytiva) using the AKTA Pure purification system. The binding buffer was 20 mM Sodium Phosphate (Na_2_HPO_4_), 0.5 M NaCl, 20 mM Imidazole, pH 7.4, and the elution buffer was 20 mM Sodium Phosphate (Na_2_HPO_4_), 0.5 M NaCl, 500 mM Imidazole, pH 7.4. The eluate was then buffer exchanged to Dulbecco’s phosphate buffered saline (DPBS) with calcium and magnesium (Gibco™ 14040133) using 7K MWCO (molecular weight cut off) Zebaspin desalting columns (Thermo Fisher). Protein aliquots were stored in -80 °C for long-term storage and at 4 °C for short-term use and storage.

### Size-exclusion chromatography

The putative oligomeric state of NA proteins was assessed via size-exclusion chromatography (SEC). Proteins were run on Superdex^®^ 200 Increase 10/300 GL column (Cytiva) equilibrated with PBS at a flow rate of 0.5 mL/min. Graphs were plotted using GraphPad Prism software.

### NA enzymatic activity assays

The enzymatic activity of NA proteins was measured using ELLA (Enzyme-linked lectin assay; large substrate fetuin Mw: 49 kDa) and MUNANA (20-(4-methylumbelliferyl)-a-D-N-acetylneuraminic acid; small fluorescent substrate Mw: 489 Da) assays.

**ELLA:** ELLA was performed as previously described ^21,24^. In brief, serially-diluted heat-inactivated sera were incubated with pre-titrated virus (NA source) or recombinant NA protein for 1 h. The dilution medium was DMEM (Gibco) supplemented with 2 mM glutamine, 10 mM HEPES, 0.1% (w/v) BSA, 100 U/mL penicillin and 100 µg/mL streptomycin. The virus or NA concentration was pre-determined by titrating to measure the lowest concentration at top plateau, ensuring at least a 5-fold signal-to-noise ratio. The mixture was transferred to a Nunc Immunoassay ELISA plate (Thermo Fisher, 439454) pre-coated overnight with 25 μg/mL fetuin (Sigma-Aldrich, F3385). The plate was incubated for 18-20 h at 37 °C with 5% (v/v) CO2 in a tissue culture incubator. The NA enzymatic activity was detected by adding HRP (Horseradish peroxidase)-conjugated peanut agglutinin (Sigma-Aldrich, L7759) at 1 μg/mL after washing the plates 3 times with PBS and then developing with 50 µL TMB (3,3ʹ,5,5ʹ-Tetramethylbenzidine) substrate (SeraCare). The enzymatic reaction was stopped after 5-10 min using 50 µL 1 M H_2_SO_4_ and the absorbance was read at 450 nm using a CLARIOstar plate reader (BMG Labtech).

**MUNANA:** The protocol from the WHO Worldwide Influenza Centre, Crick Institute was adapted. The sialidase activity of NA proteins or viruses were measured by incubation of serially-diluted NA (50 uL) in 32.5 mM MES (2-(N-morpholino)ethanesulfonic acid) buffer pH 6.5 containing 4 mM CaCl_2_ with 50 µL of 100 µM MUNANA substrate. The reaction mix was incubated for 1 h at 37 °C and stopped by adding 50 µL stop solution (0.1 M glycine, 25% ethanol, pH 10.7). Changes in fluorescence was measured at 460 nm (excitation 355 nm) using a CLARIOstar. The NA concentration giving fluorescence intensity of 40-60K units (∼100-fold signal from background) was used in NA inhibition assays. NA inhibition titres of anti-sera or mAbs were determined by incubation of serially diluted sera/mAbs with NA/viruses for 1 h before the addition of MUNANA substrate.

% NA activity was calculated as {(X—Min)/(Max—Min)}×100 where X = measurement value, Min = buffer only, Max = NA or virus alone. The 50% inhibiting titre/concentration (IC_50_) was determined using non-linear regression curve fit on GraphPad Prism. Kruskal-Wallis test was used for statistical analysis. All graphs were plotted using GraphPad Prism software.

### nanoDSF thermal unfolding

The thermal stability and unfolding of NA proteins were measured using the nanoDSF (differential scanning fluorimetry) technique on a Prometheus Panta instrument (Nanotemper). NanoDSF monitors changes in the intrinsic tryptophan or tyrosine fluorescence resulting from alterations in the 3D structure of proteins as a function of temperature. Capillaries were filled with 10 μL of NA protein at

0.4 mg/mL, placed into the sample holder and the temperature was increased from 20 to 90°C at a ramp rate of 1°C/min, with one fluorescence measurement taken per 0.1°C. The fluorescence intensity ratio (Em_350 nm_/Em_330 nm_) and its first derivative were calculated using the manufacturer’s software. Two measurements were taken for each experiment, and the means are shown. The experiment was repeated to ensure reproducibility.

### Epitope specificity analysis

A collection of monoclonal antibodies developed in-house and published in the literature was used to determine their binding specificity to NA proteins. Antibodies were serially diluted 5-fold from 10,000 to 1 ng/mL. mAbs were tested for binding to recombinant protein or NA expressed on the surface of MDCK-SIAT1 cells infected overnight with the virus. Antibody binding was detected using a secondary goat-anti human HRP antibody. The area under the curve was measured for each mAb’s titration curve and normalised as a percentage of relative binding to one of the highest binding mAb for comparison. The binding experiments were repeated, and the data from a single experiment are included. Data analysis and graphs generation was performed using GraphPad Prism.

### mi3 VLP conjugations

SpyCatcher003-mi3 virus-like particles (Lot No. 108-23-001-007) produced in *Bacillus subtilis*, were kindly provided by Ingenza Ltd. NA and VLP conjugations were done in DPBS (Dulbecco’s phosphate buffered saline) with Calcium and Magnesium (Gibco™ 14040133) at various molar ratios. The conjugation was analysed using reduced SDS-PAGE. The conjugates were confirmed to have enzymatic activities and binding of mAbs binding equivalent to the unconjugated proteins.

### Mice immunisation and virus challenge

Animal experiments were conducted in compliance with the UK Animals (Scientific Procedures) Act Project License (PP9362617), following the principles of the 3Rs (Replacement, Reduction and Refinement). Female BALB/c OlaHsd or DBA/2 OlaHsd mice (7-8 weeks old at the start of the experiment) were obtained from Envigo and housed in individually vented cages in a specialised unit for infectious diseases. The mice were housed in accordance with the UK Home Office ethical and welfare guidelines, fed standard chow, and had access to water *ad libitum*. Mice were anesthetized with isoflurane (Abbott) and immunised via intramuscular injections with two doses of 0.5 µg of NA-VLP immunogens adjuvanted with AddaVax^TM^ 1:1 vol/vol, administered 3 weeks apart. Serum samples were collected 3 weeks after the booster dose: intermediate samples were obtained via tail vein and terminal samples via cardiac puncture of euthanised mice. Whole blood collected in microtainer SST tubes (BD) was allowed to clot at room temperature for 1-2 h before centrifugation at 10,000 *g* for 10 min. The clarified sera were transferred to fresh tubes and stored at -20 °C.

For virus challenge experiments, 50 uL virus (passaged and titrated in the laboratory) was administered intranasally to anesthetised mice. Virus dose was 10^4^ TCID_50_ (tissue culture infectious dose 50%) [200 LD_50_ (lethal dose 50%)] of X-179A (H1N1 A/California/07/2009) in DBA/2 mice and 10^4^ TCID_50_ (1000 LD_50_) of H1N1 A/PR/8/1934 in BALB/c mice. The weight and clinical signs of mice were monitored regularly over a study period of 2-3 weeks until they either reached their endpoint (≤80% of their initial weight) or recovered to their initial weight. Mice reaching the endpoint were humanely euthanised. Kaplan-Meier survival analysis with the Logrank Mantel-Cox test was used for comparison.

### Sequence Alignment

Neuraminidase sequences were obtained from PDB files where referred, or from Global Initiative on Sharing All Influenza Data (GISAID). Amino acid sequences of the neuraminidase were aligned using Muscle alignment in Geneious Prime, and all alignment figures were generated using Geneious Prime.

### Crystallization, data collection and structure determination

In each well of 96-well sitting drop plates (Greiner), 100 nL of purified neuraminidase was combined with 100 nL of precipitant. The concentrations used for crystallisation were 4.6 mg/mL for mSN1, 5 mg/mL for N1/09 hybrid and 4.5 mg/mL for N1/19 hybrid. For crystal formation, the mixtures were equilibrated against 90 μL of precipitant at 20.5 °C ^43^.

Crystals of msN1 were obtained using the Morpheus B2 precipitant [0.09M halogens (NaFl, NaBr, NaI), 0.1 M imidazole/MES pH 6.5, 30% ethylene glycol/PEG (polyethylene glycol) 8K; Hampton Research]. The N1/09 hybrid crystallized using precipitant H4 of the Pentaerytritol screen (25% pentaerytritol ethoxylate 797, 0.1 M MES, 0.05 M MgCl2, pH 6.5; Jena Bioscience) and the N1/19 hybrid using PGA (poly gamma glutamic acid)-screen condition G2 (5% PGA-low molecular weight, 30% PEG550 MME (monomethyl ether), 0.1 M Tris pH 7.8; Molecular Dimensions). Glycerol was added to 25% (v/v) for cryoprotection of the N1/19 hybrid crystals. Diamond beamline I24 (Harwell, UK) was used for diffraction data collection at 100K. Data processing used the Xia2 programme suite ^44^ and, for the N1/19 hybrid, autoPROC and STARANISO ^45^. The molecular replacement program PHASER ^46^ was used for structure elucidation. Model building was done using COOT ^47^ and model refinement using Phenix^48^. Data-collection and refinement statistics are provided in Supplementary Table 3. Figures were prepared using UCSF ChimeraX ^22^.

## Data Deposition

The atomic coordinates and structure factors for the mSN1, N1/09 hybrid and N1/19 hybrid structures have been deposited in the Protein Data Bank with the accession codes 9GQX, 9GQT and 9GQQ, respectively.

## Acknowledgements

We thank Ingenza Ltd for providing SpyCatcher003-mi3 particles, and Dr Rodney Daniels and Dr John McCauley for providing influenza viruses. Thanks to Max Quastel, Dr Jack Tan, Lisa Schimanski, and Diana Melnyk for laboratory assistance. The authors would also like to thank Diamond Light Source (Harwell, UK) for beamtime (proposal mx28534), and the I24 beamline staff for assistance with data collection. This work was supported by the Chinese Academy of Medical Sciences (CAMS) Innovation Fund for Medical Science (CIFMS), China (grant number: 2018-12M-2-002) to P.R. and A.T., and the Townsend-Jeantet Prize Charitable Trust Charity No. 1011770. The work of monoclonal antibody development was supported by the National Science and Technology Council of Taiwan (NSTC 111-2628-B-002-053-, 112-2314-B-002-099-MY3) to K.-Y.A.H. The structural work was funded by the Medical Research Council MR/S007555/1 to T.A.B.. The Centre for Human Genetics was supported by the Wellcome grant 203141/Z/16/Z.

## Author Contributions

P.R. and A.T. designed the research; P.R., L.W., and G.C.P. performed the research; K.-Y.A.H. contributed antibodies; P.R., L.W., G.C.P., T.A.B., D.S., and A.T. analysed the data; P.R., A.T., K.-Y.A.H., and T.A.B. acquired the funding; P.R. and A.T. wrote the manuscript with input from all authors.

## Competing Interests

A.T. and P.R. are inventor on patents filed on Influenza Neuraminidase loop-based design (Pending).

M.R.H. is an inventor on patents filed on spontaneous amide bond formation (EP2534484 and UK Intellectual Property Office 1706430.4) and a SpyBiotech co-founder and shareholder.

## Notes

### Summary of Updates

1. Abstract: Influenza neuraminidase has been changed to Influenza virus neuraminidase, and conferred protected has been corrected to protected. 2. et al. are italicised 3. The previous lines 91-96 have been moved to the second Results section (new lines 104-108) to improve flow and maintain the order in which figures are mentioned. 4. Line 151: Manuscript in preparation, has been changed to Personal communication with Dr. Yan Wu, Capital Medical University. 5. All abbreviations have been defined. 6. Figure references have been added within the Results section, under Murine Challenge Studies. 7. A sentence has been added to the Results section, under Loop transfer between two distant N1 NAs: ......., to clarify the rationale for performing the loop transfer on PR8 NA. 8. Line 272: The figure reference has been corrected. 9. A sentence has been added within the Discussion section (The distribution of epitopes on neuraminidase, lines 369-371) to emphasize that the majority of antibody epitopes (∽90%) are located within loops L01 and L23, based on a compilation of epitopes from published NA mAbs and polyclonal sera (via escape mutagenesis and NA-Fab crystal structures; Supplementary Figures 1 and 2). L01 and L23 primarily form the top surface but also extend to regions described in some literature as the side, lateral, underside, and edges of NA. 10. Figure 1 has been updated to reflect this point. 11. An additional reference (Ref 38) has been added to line 369. 12. Figure 4 and its description have been updated to improve clarity and informativeness, following the reviewers comments. 13. Figures 3a and 6b have been updated to include EC50 and area under the curve values for NA enzymatic titration curves. 14. An additional panel (Figure 7d) has been added to Supplementary Figure 7 to demonstrate that binding titres are equivalent to ELLA IC50 titres. 15. The Supplementary Figure 5 description has been updated for clarity.

