## Supplemental Data for "Structure-Guided Loop Grafting Improves Expression and Stability of Influenza Neuraminidase for Vaccine Development"

Pramila Rijal[1,2], Leiyan Wei[1,2], Guido C Paesen[3], David I Stuart[1,3], Mark R Howarth[4], Kuan-Ying A Huang[5], Thomas A Bowden[3], Alain RM Townsend[1,2]

**Supplementary Table 1. Recombinant NA expression in various cell expression systems.**

| *Literature* | Subtype (NA) | Virus type | Expression System | Yield |
|---|---|---|---|---|
| *Prevato 2015* [1] | N1 | H1N1/09 | Expi293F | **30 mg/L** |
| *Ellis 2022* [2] | N1, N2, N8 | H1N1/09, H1N1/15, H3N2/05, H10N8/13 | Expi293F | **up to 30 mg/L** |
| *Ecker 2020* [3] | N1, N2 | H1N1/09 H1N1/18, H3N2/13, H3N2/14, H3N2/16 | Expi293F | **up to 28 mg/L** |
| *Prevato 2015* [1] | N1 | H5N1 | Expi293F | **6 mg/L** |
| *Martinet 1997* [4] | N2 | H3N2/1975 | *P Pastoris* | **2.5–3 mg/L** |
| *Woude 2020* [5] | N2 | H3N2 | 293T | **2.5 mg/L** |
| *Subathra 2014* [6] | N1 | H1N1 | *P Pastoris* | **2 mg/L** |
| *Schmidt 2011* [7] | N1 | H1N1 | Sf21 | **0.5-1.8 mg/L** |
| *Nivitchanyong 2011* [8] | N1 | H5N1 | 293-F | **0.3-0.7 mg/L** |
| *Margine 2013* [9] | N9 | H7N9/2013 | Sf9 | **0.2-0.7 mg/L** |
| *Liu 2015* [10] | N1 | H1N1/09, H5N1 | Sf9 | **0.25-0.5 mg/L** |

**Supplementary Table 2.** List of loop annotations in Varghese et al. [11] used as a reference for creating NA hybrids in this paper. Loops in mSN1, N1/09, N1/19 and PR8N1 are listed. Text in orange colour where N1 differs from the original N2 numbering system. Residues in the loops that differ from mSN1 sequence are marked in red. ('aa' denotes amino acids).

| Loops | N2 numbering | N2 NA Loops (Varghese et al 1983) | N1 numbering | mSN1 Loops | N1/09 Loops | N1/19 Loops | PR8 N1 Loops |
|---|---|---|---|---|---|---|---|
| B1L01 | 107-119 | RLSAGGDIWVTRE | 107-119 | RIGSKGDVFVIRE | RIGSKGDVFVIRE | RIGSKGDVFVIRE | RIGSKGDVFVIRE |
| B1L23 | 135-156 | GQGTTLDNKHSNDTVH DRIPHR | 135-156 | TQGALLNDKHSNGTVKDR SPYR | TQGALLNDKHSNGTIKDRS PYR | TQGALLNDKHSNGTIKDRS PYR | TQGALLNDRHSNGTVKDR SPYR |
| B2L01 | 175-177 | CIA | 176-178 | SVA | SVA | SVA | SVA |
| B2L23 | 195-199 | TGDDK | 196-199 | SGPD | SGPD | SGPD | SGPD |
| B3L01 | 218-227 | WSQNILRTQE | 219-228 | WRNNILRTQE | WRNNILRTQE | WRNKILRTQE | WRKKILRTQE |
| B3L23 | 243-250 | DGSASGRA | 244-251 | DGPSNGQA | DGPSNGQA | DGPSDGQA | DGPSDGLA |
| B4L01 | 269-277 | AGSAQHVEE | 269-278 | LNAPNYHYEE | MNAPNYHYEE | MKAPNYHYEE | LNAPNSHYEE |
| B4L23 | 292-295 | RDNW | 293-296 | RDNW | RDNW | RDNW | RDNW |
| B5L01 | 315-350 | SYVCSGLVGDTPRNDD RSSNSNCRDPNNERGT | 314-347 | IGYICSGVFGDNPRPNDGT GSCSPMSSNGAYGVK | IGYICSGIFGDNPRPNDKTG SCGPVSSNGANGVK | MGYICSGVFGDNPRPNDK TGSCGPVSSNGANGVK | IGYICSGVFGDNPRPKDGT GSCGPVYVDGANGVK |
| B5L23 | 367-371 | SKDLR | 364-369 | STSSRS | SISSRN | SISSRK | SHSSRH |
| B6L01 | 399-403 | DSDNR | 395-399 | EITDW | GINEW | GINEW | AMTDW |
| B6L23 | 429-437 | GRKQETRVW | 430-437 | RPKENTIW | RPKENTIW | RPEENTIW | GRPKEKTIW |
| Comments | | | | | 12 aa changes in 5 Loops | 16 aa changes in 8 Loops | 18 aa changes in 8 Loops |

**Supplementary Table 3. Crystallographic data collection and refinement statistics.**

| DATA COLLECTION | mSN1 | N1/09 | N1/19 |
|---|---|---|---|
| **Beamline** | DLS I24 | DLS I24 | DLS I24 |
| **Wavelength (Å)** | 0.6199 | 0.6199 | 0.6199 |
| **Space Group** | *P* 4 | *P* 4 | $C\,2\,2\,2_1$ |
| **Cell Dimensions** | | | |
| a, b, c (Å) | 92.2, 92.2, 147.9 | 92.1, 92.1, 146.8 | 115.5, 234.4, 115.7 |
| $\alpha$, β, $\gamma$ (°) | 90, 90, 90 | 90, 90, 90 | 90, 90, 90 |
| **Resolution range (Å)** | 147.90-1.98 [2.01-1.98] | 41.18-1.94 [1.97-1.94] | 82.31-1.78 [1.98-1.78] |
| **Rmerge** | 0.766 [2.784] | 0.792 [3.109] | 0.367 [1.705] |
| **I/σ (I)** | 2.8 [0.7] | 4.1 [0.7] | 5.4 [1.6] |
| **$CC_{1/2}$** | 0.976 [0.342] | 0.983 [0.282] | 0.988 [0.564] |
| **Completeness (%)** | 100 [99.8] | 99.9 [98.3] | 94.2 [65.2]* |
| **Multiplicity** | 14.2 [14.1] | 13.9 [10.9] | 13.4 [12.9] |
| **REFINEMENT** | | | |
| **Resolution (Å)** | 92.17-1.98 | 41.18 -1.94 | 64.69-1.78 |
| **No. reflections** | 83,874 | 89,971 | 58,510 |
| **$R_{work}/R_{free}$** | 0.211/0.238 | 0.183/0.213 | 0.233/0.267 |
| **No. atoms** | | | |
| protein | 6,296 | 6,279 | 6,164 |
| ligands | 127 | 102 | 116 |
| solvent | 619 | 695 | 474 |
| **Average B-factors** | | | |
| protein | 29.3 | 24.2 | 18.23 |
| ligand | 65.4 | 66.3 | 60.12 |
| solvent | 35.8 | 33,6 | 25.0 |
| **Ramachandran (%)** | | | |
| favoured | 96.21 | 96.32 | 96.22 |
| allowed | 3.55 | 3.68 | 3.58 |
| outlier | 0.24 | 0 | 0.25 |
| **RMS** | | | |
| bond lengths (Å) | 0.003 | 0.013 | 0.004 |
| bond angles (°) | 0.67 | 1.16 | 0.71 |

The values between brackets are for the highest-resolution shell
*Ellipsoidal completeness as determined by STARANISO [12]

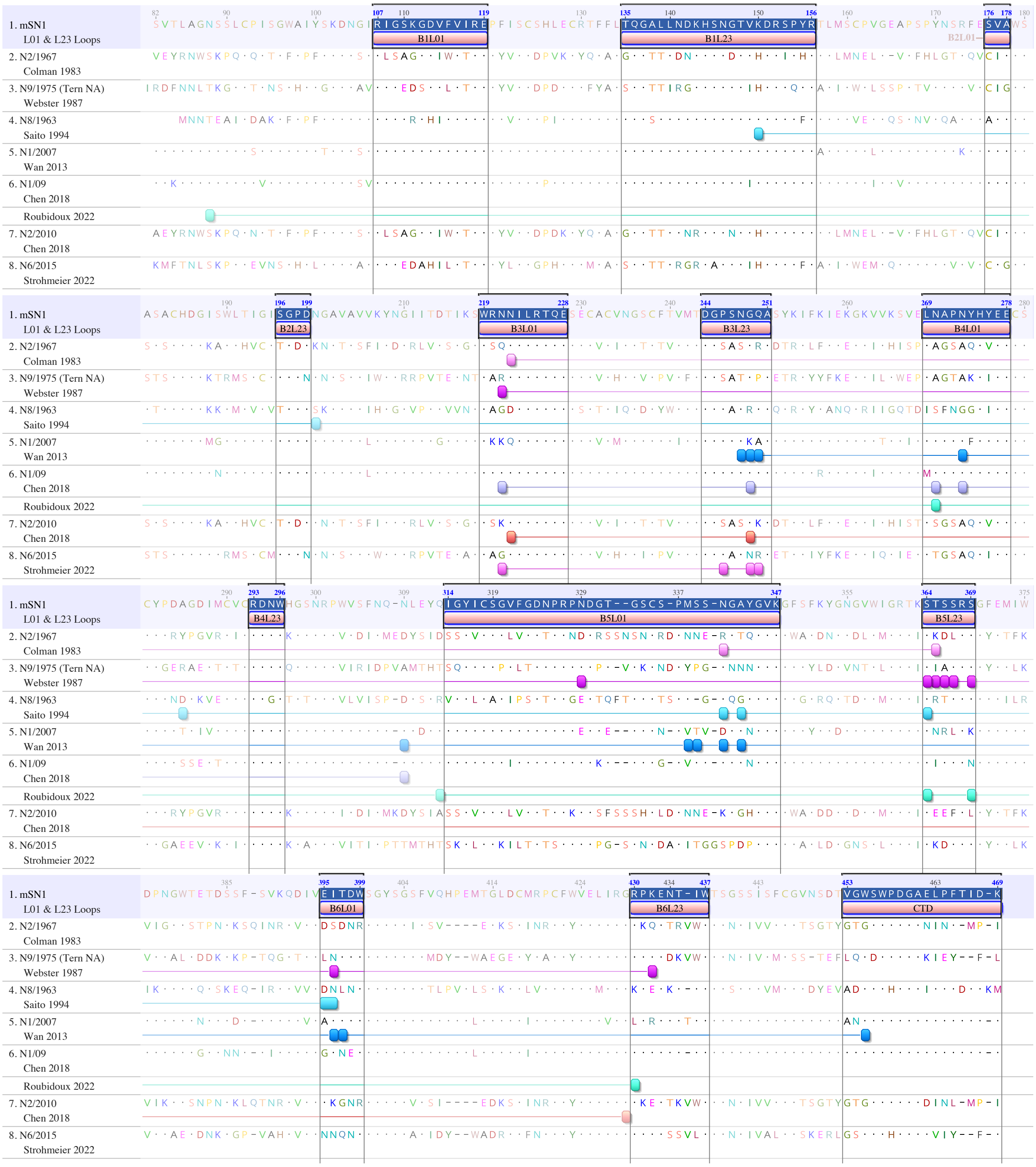

**Supplementary Figure 1**. Viral neuraminidases selected for resistance to various monoclonal antibodies and sera with the positions of substitutions shown from seven published studies (N1 numbering with H5N1 A/mute swan/England/053054/2021 (mSN1 as reference). The antibody epitopes from each study are shown in a different coloured blocks with the sequence of the original NA used in these studies. Twelve L01 and L23 loops and C-terminal domain are highlighted. The figure shows 45 examples of mAb selection experiments. Fourteen amino acid positions were selected repeatedly (13 in loops 01 and 23, 1 at 309 on the underside of the head at B4L34) by different antibodies so that in total 31 amino acid residue positions have been identified as sites for selection of resistant viruses. Among these 31 sites, 25 are in L01 and L23 or CTD as defined, 3 are immediately adjacent to L01 and L23 and could be considered as part of the top loops in future designs, and 3 sites are in underside positions: at 88 between stem and head, 285 on B4L12, and 309 on B4L34. The mSN1 sequence and the loop assignments are based on Varghese et al. 1983 and 1992 as reference [11,13]. This shows that ~90% of sites for selection by monoclonal and serum antibodies of resistant viruses reside within the L01, L23 and CTD. Figure was generated using Geneious Prime. References: [14,15,16,17,18,19,20].

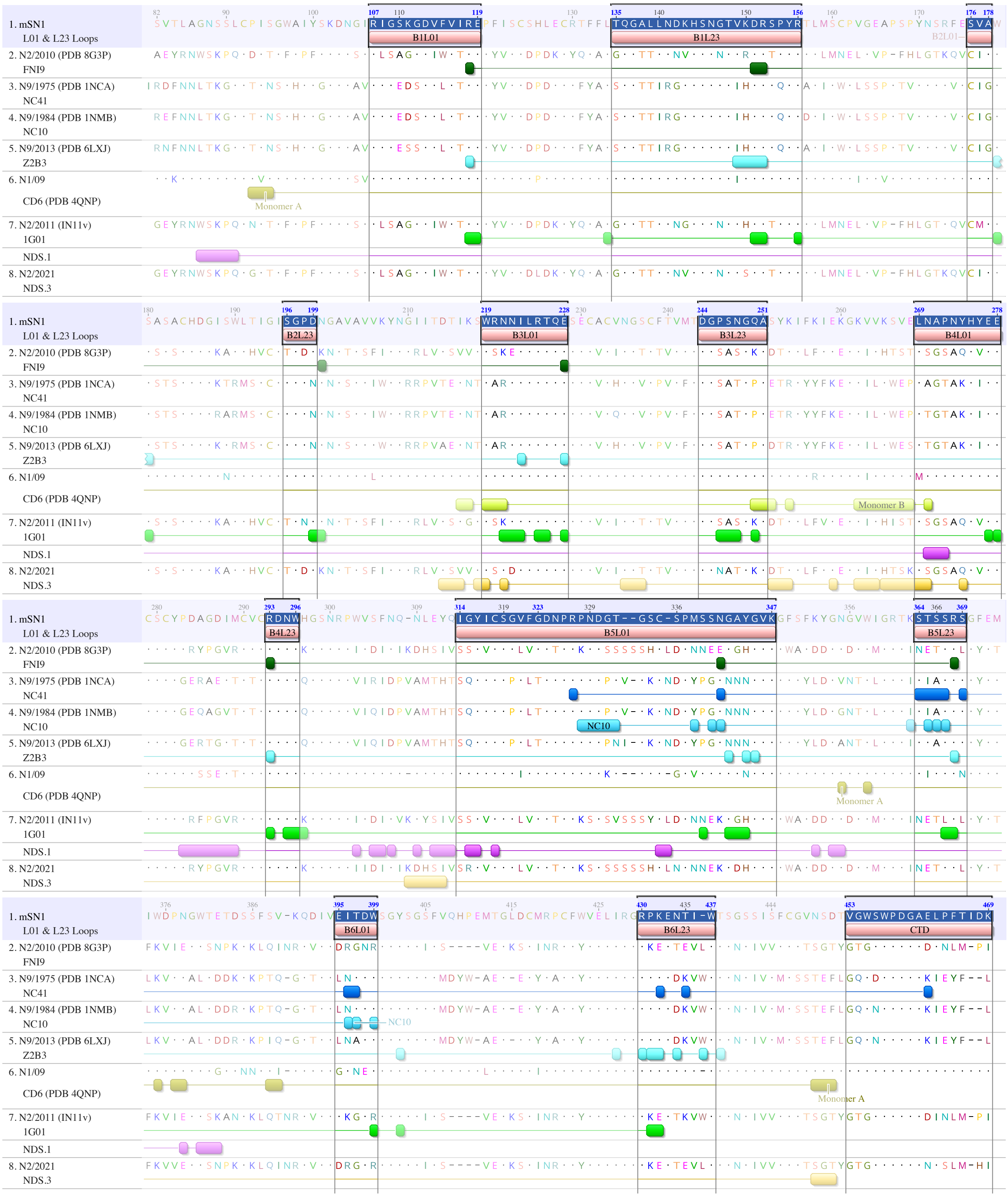


**Supplementary Figure 2**. Sequence alignment of various neuraminidases showing the contact residues for various mAbs, as defined by cryo-EM or crystal structures. mAb binding epitopes from each study are shown in a different coloured blocks with the sequence of the original NA used in these studies. The msN1 sequence is included as a reference for sequence alignment, and twelve L12 and L23 loops and the C-terminal domain are highlighted. mAbs NDS.1 and NDS.3 are described as antibodies towards underside loops. However, they form some interactions with L01 and L23 as well. This shows that the majority of key antigenic sites are located within the surface Loops 01 and 23, with some on

the underside of the NA head, confirming the data from selections of resistant viruses by NA specific MAbs. Figure was generated using Geneious Prime.
References: FNI9 [21], NC41 [22–24], NC10 [25,26], Z2B3 [27], CD6 [28], 1G01 [29,30], NDS.1 and NDS.3 [29].

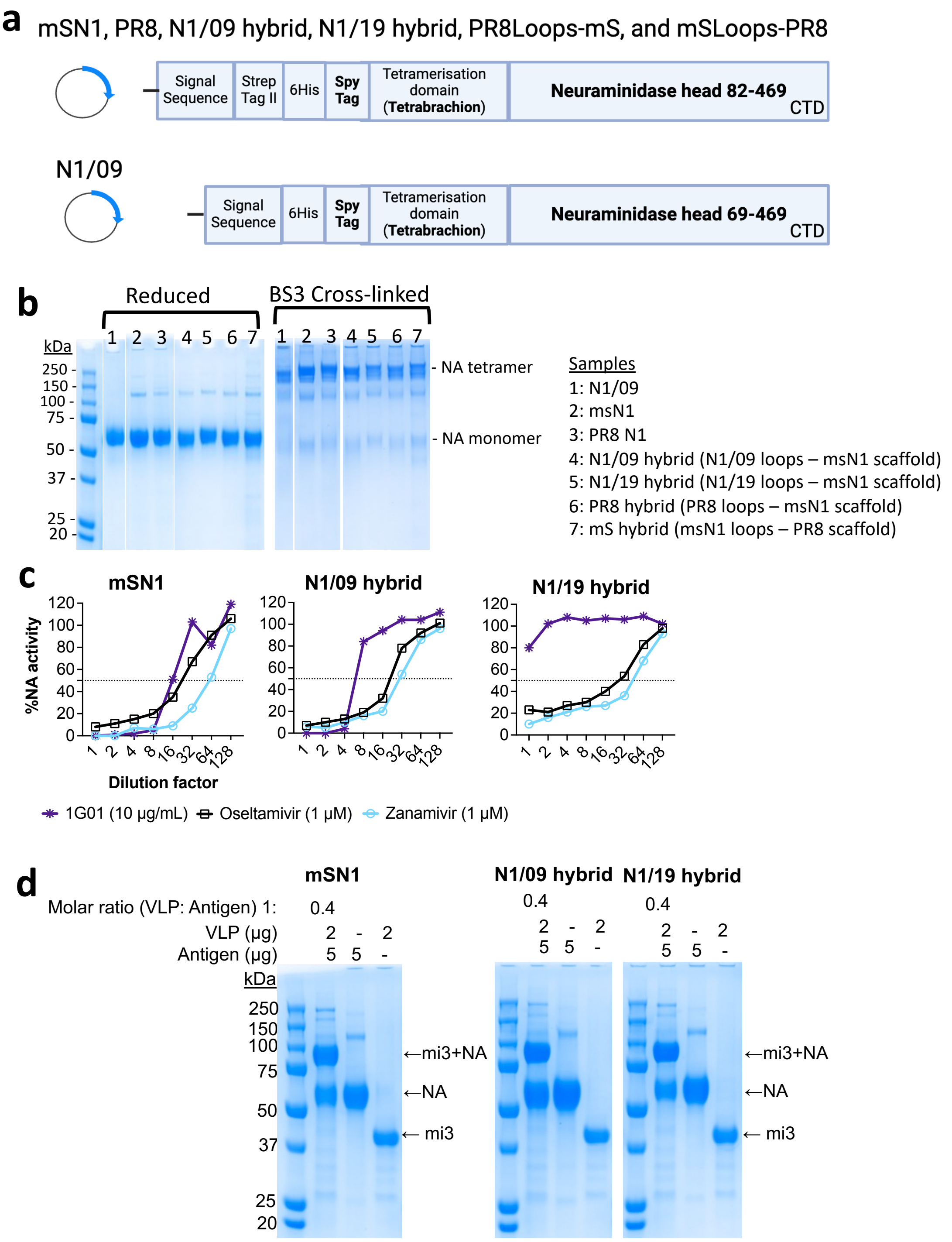


**Supplementary Figure 3. a) Gene constructs for NA proteins.** All the constructs are the same except for N1/09 which uses the Ig Kappa signal sequence, contains extra residues 69-81 in the NA head and has no Strep tag II. **b) NA and hybrid proteins were expressed as tetramers.** Purified proteins on SDS-PAGE with Coomassie staining, in reducing conditions and cross-linked before loading on the gel. 2 µg of NA protein was incubated with 6 mM BS3 cross-linking reagent for 30 min and the reaction was stopped by adding 1M Tris-HCl pH 8.0. Proteins were separated on reducing 4-12% Bis-Tris SDS-PAGE and 1x MES SDS buffer. **c) Inhibition of enzyme activity of NA hybrid proteins by small molecule inhibitors and mAbs.** Oseltamivir and Zanamivir inhibit the function of mSN1, N1/09, and N1/19 hybrid proteins suggesting the active site is intact and functional. mAb 1G01 [30] inhibited mSN1 and N1/09 hybrid suggesting the cross-reactive epitopes recognised by 1G01 have been preserved. 1G01 fails to inhibit N1/19 due to substitution at N222K (see figure 2 alignment) and this has been confirmed by the loss of inhibition of NA activity of N1/19 hybrid. **d) Coupling of NA to mi3 virus-like particles (VLP) using SpyCatcher technology.** SDS-PAGE showing the NA+mi3, NA and mi3. Two µg mi3 with SpyCatcher003 tag covalently linked to 5 µg NA (molar ratio 1:0.4). These NA-VLPs were used for mouse immunisations.

**a**

| Loops | Varghese 1983 Annotations | | Varghese 1991 Annotations | |
|---|---|---|---|---|
| | *N2 numbering* | *Sequence* | *N2 numbering* | *Sequence* |
| B1L01 | 107-119 | RLSAGGDIWVTRE | 104-116 | NSIRLSAGGDIWV |
| B1L23 | 135-156 | GQGTTLDNKHSNDTVHDRIPHR | 137-156 | GTTLDNKHSNDTVHDRIPHR |
| B2L01 | 175-177 | CIA | 176-177 | IA |
| B2L23 | 195-199 | TGDDK | 196-200 | GDDKN |
| B3L01 | 218-227 | WSQNILRTQE | 218-220 | WSQ |
| B3L23 | 243-250 | DGSASGRA | 246-248 | ASG |
| B4L01 | 269-277 | AGSAQHVEE | 269-275 | AGSAQHV |
| B4L23 | 292-295 | RDNW | 293-299 | DNWKGSN |
| B5L01 | 315-350 | SYVCSGLVGDTPRNDDRSSNSNCRD PNNERGTQGVK | 318-351 | CSGLVGDTPRNDDRSSNSNCRDPN NERGTQGVKG |
| B5L23 | 367-371 | SKDLR | 365-373 | TISKDLRSG |
| B6L01 | 399-403 | DSDNR | 399-405 | DSDNRSG |
| B6L23 | 429-437 | GRKQETRVW | 431-437 | KQETRVW |

**b** **c** **d** **e**

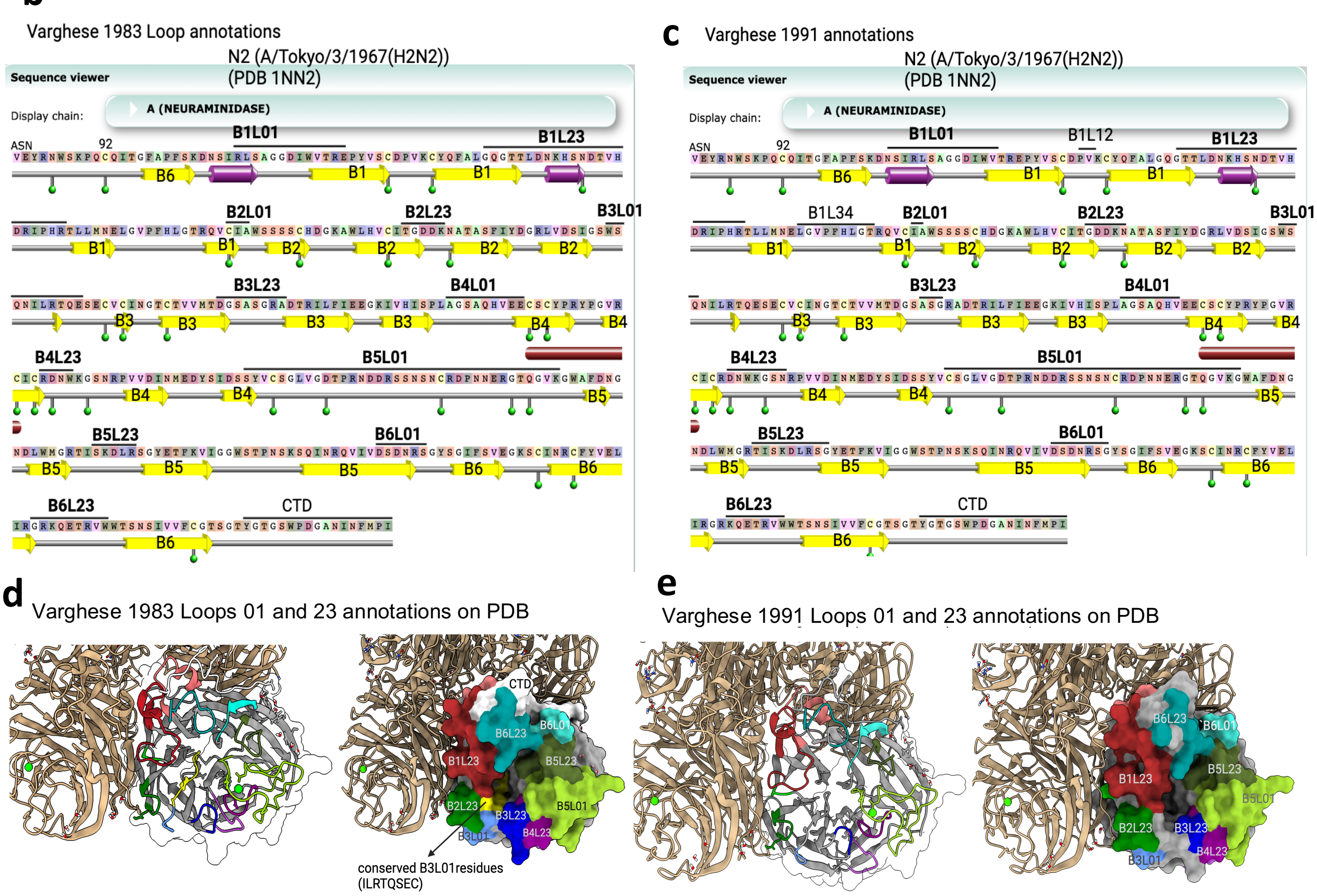


**Supplementary Figure 4. a,b,c,** Loops annotations in Varghese et al 1983 and 1991 [13,14] on PDB 1NN2 N2 NA sequence. Figures were downloaded and adapted from pdbj.org (b,c). **d,e,** Loops were visualised and coloured on PDB 1NN2 structure using Chimera X1.4.

**a**

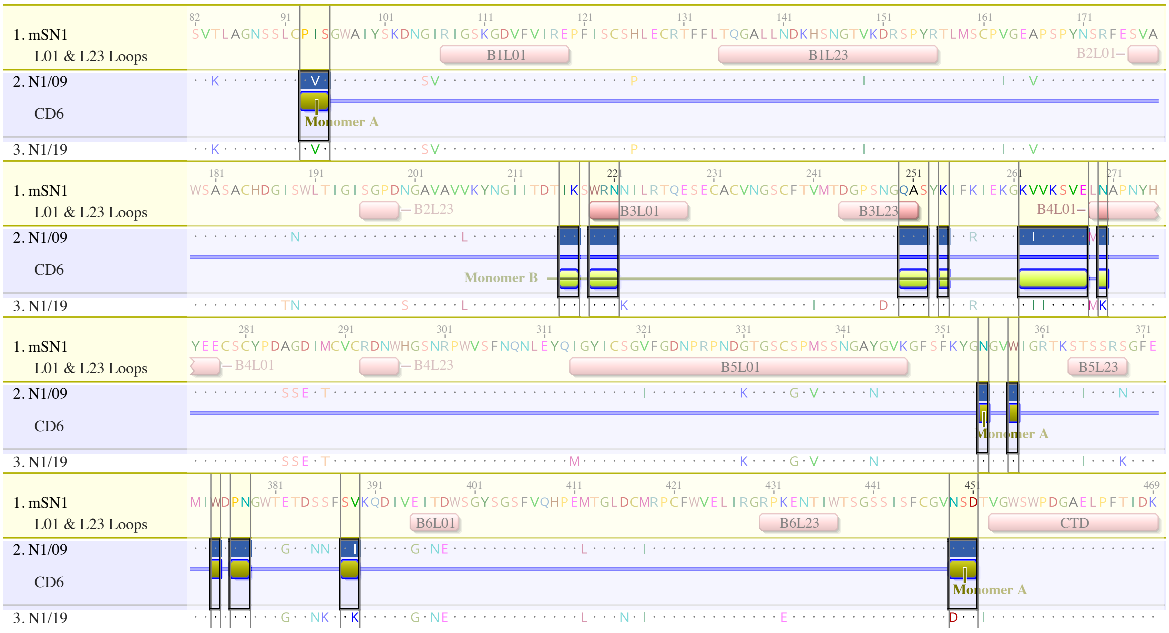


**b**

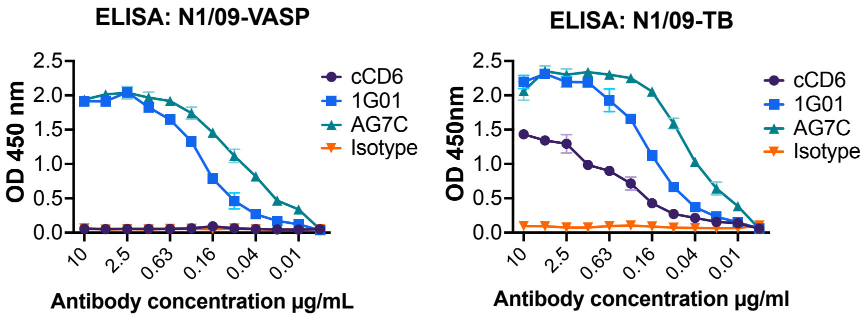


**Supplementary Figure 5. a, Binding by mAb CD6 is predominantly scaffold dependent and occurs across two protomers.** Epitope recognised by mAb CD6 on H1N1/09 (PDB 4QNP). mAb CD6 [28] forms interactions with two monomers, indicated with olive green and light green. msN1 sequence is used as a reference showing Loops 01 and 23. Interactions with monomer A (olive green) are all on the NA scaffold, whereas interactions with monomer B (light green) are on the top surface Loops and scaffold. CD6 binds to N1/09 and mSN1 but not to N1/19. N1/19 escape from mAb CD6 is likely due to one or more of the following substitutions within the epitope - L269M (B4L01), N270K (B4L01), V389K (B5S4), and N449D (B6L34). However, the N1/19 hybrid that shares two of these substitutions L269M (B4L01) and N270K (B4L01) retained binding, which suggests that loss of binding to N1/19 was due to V389K (B5S4) and or N449D (B6L34). **b, Recognition of N1/09 NA protein expressed as tetramers with VASP or tetrabrachion tetramerisation domains**. Note that CD6 did not bind to N1/09-VASP and but did bind to N1/09-TB relatively weakly compared to mAbs AG7C [31] and 1G01 [30] that (unlike CD6) bind within a single monomer.

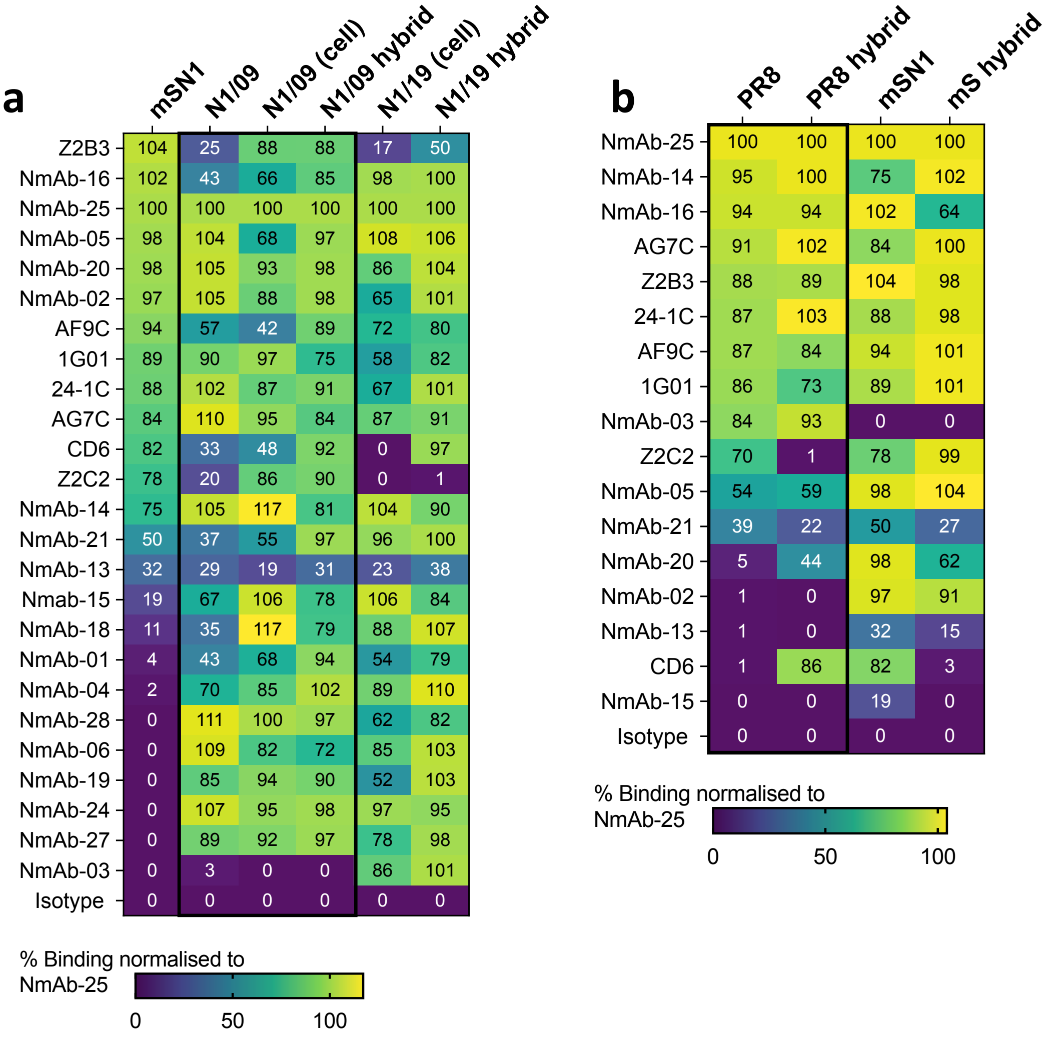


**Supplementary Figure 6.** Binding titration of mAbs against recombinant soluble proteins or NA expressed on the surface of virus infected cells. **a)** Twenty-five anti-N1 mAbs were titrated for binding against mSN1, N1/09 and N1/19 and their loop transferred hybrid variants. **b)** Binding titration of eighteen mAbs that bind either PR8 N1 or mSN1 on wild type sequence proteins and their loop-grafted hybrid proteins. Area under curve (AUC) was calculated and normalised against one of the strongest binders to rank the order.

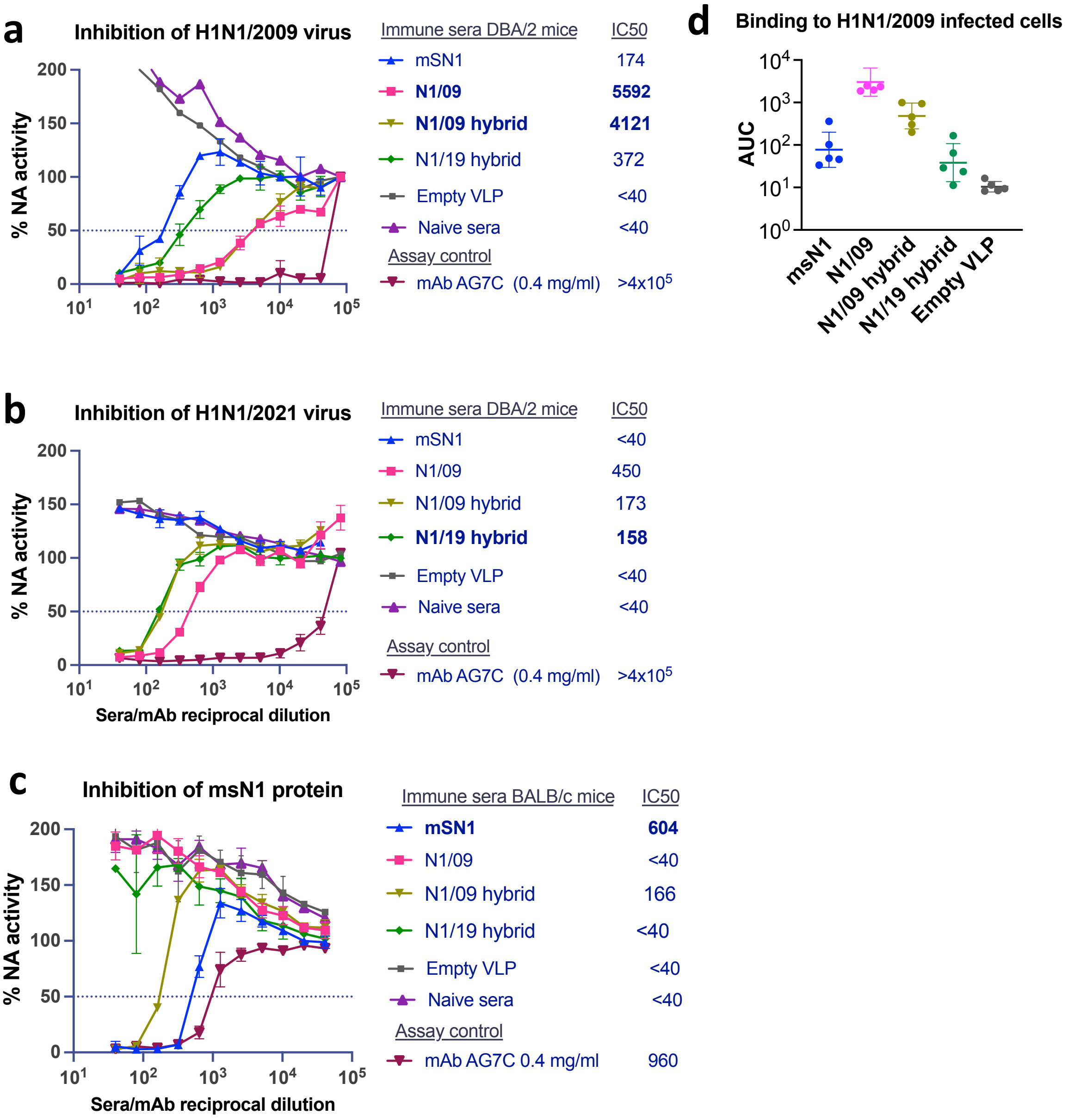


**Supplementary Figure 7. a-c)** Pooled sera from immunised mice (n=6) in Figure 5a (BALB/c mice) and Figure 5b (DBA/2 mice) were titrated in ELLA assay. Naïve sera and sera from mice immunised with empty VLP were used as negative controls. mAb AG7C was used as a positive assay control. H1N1/2009 virus = X-179A (A/California/07/2009); H1N1/2021 = A/Sydney/5/2021 (Differs from N1/19 only by V453M in the C-terminal domain), mSN1 = NA from H5N1 A/mute swan/England/053054/2021. d) Binding of sera antibodies to H1N1/2009 infected cells, for samples related to Figure 5b and Supplementary Figure 7a are compared. Similar to the ELLA titres, the binding titres for msN1 are lower, indicating that the protective response must be from the ELLA inhibiting antibodies. AUC: Area under curve of the titration curve.

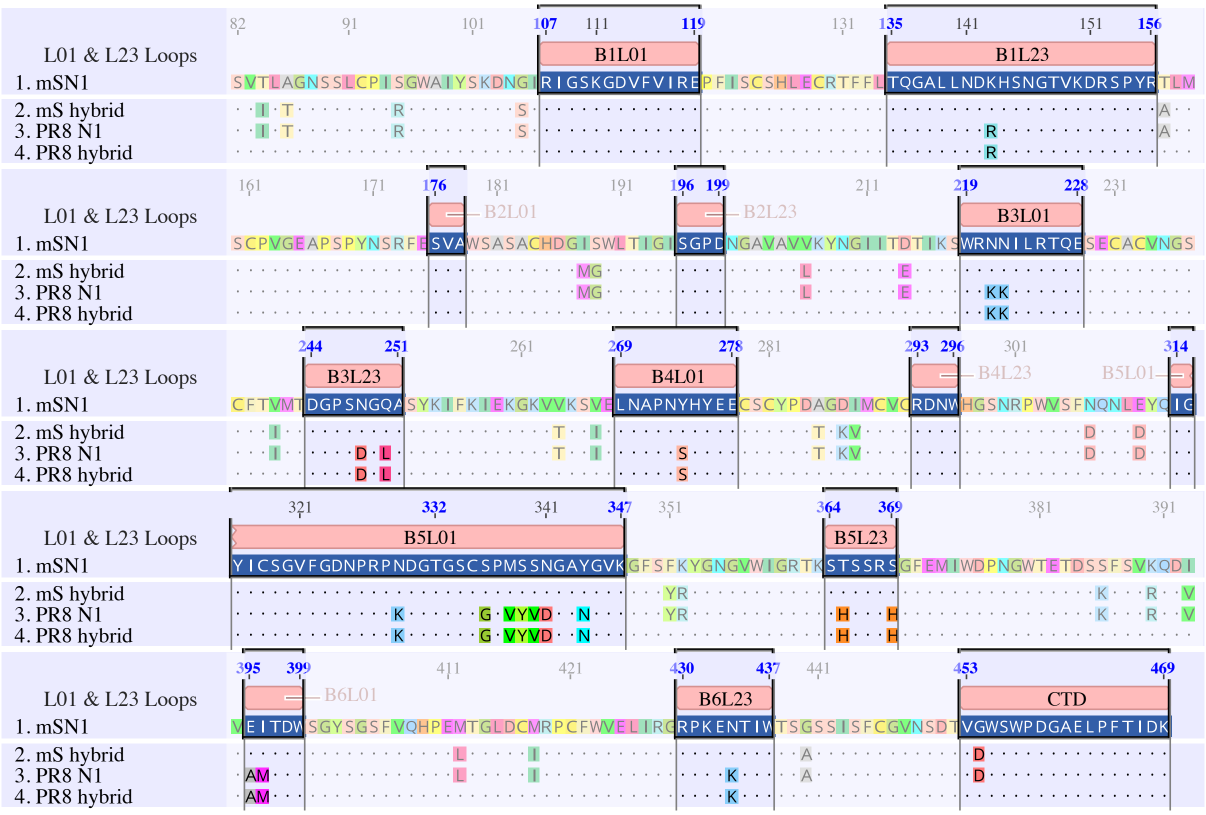


**Supplementary Figure 8. Loop transfer between two distant N1 NAs: H5N1 A/mute swan/England/053054/2021 (mS) and H1N1 A/PR/8/1934 (PR8).**

Twelve Loops 01 and 23 in a NA head of a monomer are highlighted within the sequence alignment of mSN1, PR8 N1 and loops-exchanged NA hybrid proteins. There is a residue difference at position 454 in C-terminal domain (CTD) between mSN1 and PR8N1. CTD appears at the monomeric interface and can form a part of antigenic surface. However, this residue appears not to be solvent exposed and hence was not considered for transfer. Figure was created using Geneious Prime. Refer to Figure 6.

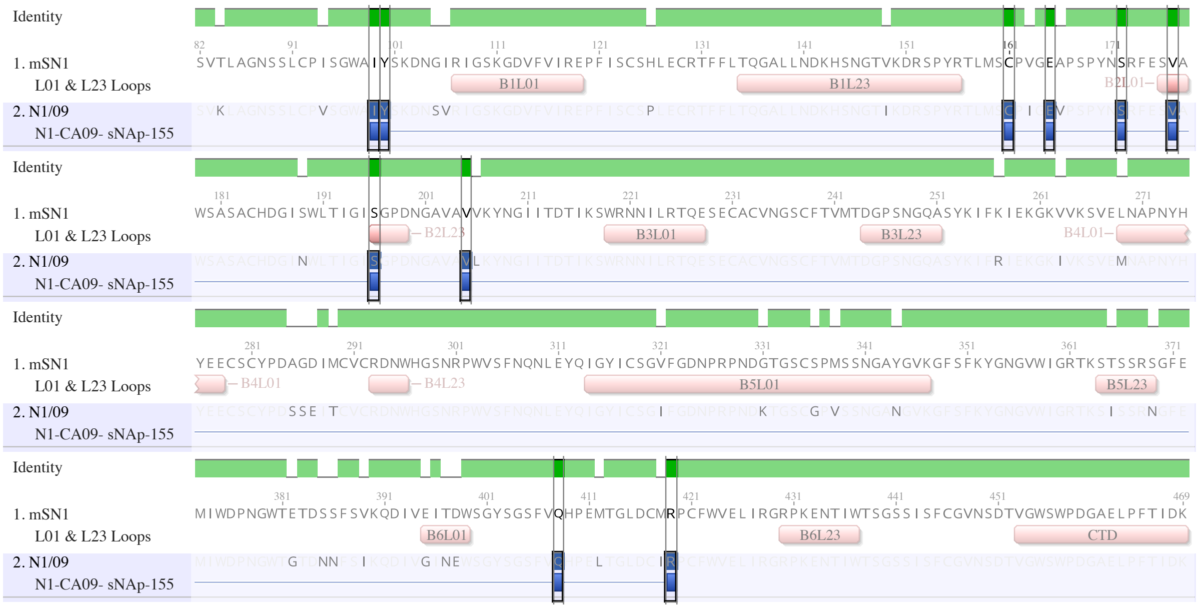

**Supplementary Figure 9. Comparison with the optimised N1 sequence by Ellis *et al.*** Ten residues mutated in N1/09 (construct N1-CA09-sNAp-155 with VASP tetramerisation domain) for favouring closed conformation by Ellis et al. are highlighted. All ten residues are unaltered in our msN1 and N1/09 hybrid proteins. Figures were created using Geneious Prime.

**Amino acid sequences of NA gene constructs**

H7 HA Signal Sequence: *MNTQILVFALIAIIPTNADKI*
IgKappa Signal Sequence: *METDTLLLWVLLLWVPGSTGD*
Strep Tag II: SAWSHPQFEK
SpyTag: AHIVMVDAYKPTK
6His Purification Tag: HHHHHH
Tetrabrachion tetramerisation domain:
IINETADDIVYRLTVIIDDRYESLKNLITLRADRLEMIINDNVSTILA
NA head sequence is coloured blue.

>mSN1 (A/mute swan/England/053054/2021)
*MNTQILVFALIAIIPTNADKI*SAWSHPQFEKGGGGSHHHHHHSSSGSGAHIVMVDAYKPTKGGSGGGG
SIINETADDIVYRLTVIIDDRYESLKNLITLRADRLEMIINDNVSTILAGGSGTGSVTLAGNSSLCPI
SGWAIYSKDNGIRIGSKGDVFVIREPFISCSHLECRTFFLTQGALLNDKHSNGTVKDRSPYRTLMSCP
VGEAPSPYNSRFESVAWSASACHDGISWLTIGISGPDNGAVAVVKYNGIITDTIKSWRNNILRTQESE
CACVNGSCFTVMTDGPSNGQASYKIFKIEKGKVVKSVELNAPNYHYEECSCYPDAGDIMCVCRDNWHG
SNRPWVSFNQNLEYQIGYICSGVFGDNPRPNDGTGSCSPMSSNGAYGVKGFSFKYGNGVWIGRTKSTS
SRSGFEMIWDPNGWTETDSSFSVKQDIVEITDWSGYSGSFVQHPEMTGLDCMRPCFWVELIRGRPKEN
TIWTSGSSISFCGVNSDTVGWSWPDGAELPFTIDK*

>N1/09 (A/California/07/2009)
*METDTLLLWVLLLWVPGSTGD*HHHHHHGSGAHIVMVDAYKPTKGGGSIINETADDIVYRLTVIIDDRY
ESLKNLITLRADRLEMIINDNVSTILAGGSGTGISNTNFAAGQSVVSVKLAGNSSLCPVSGWAIYSKD
NSVRIGSKGDVFVIREPFISCSPLECRTFFLTQGALLNDKHSNGTIKDRSPYRTLMSCPIGEVPSPYN
SRFESVAWSASACHDGINWLTIGISGPDNGAVAVLKYNGIITDTIKSWRNNILRTQESECACVNGSCF
TVMTDGPSNGQASYKIFRIEKGKIVKSVEMNAPNYHYEECSCYPDSSEITCVCRDNWHGSNRPWVSFN
QNLEYQIGYICSGIFGDNPRPNDKTGSCGPVSSNGANGVKGFSFKYGNGVWIGRTKSISSRNGFEMIW
DPNGWTGTDNNFSIKQDIVGINEWSGYSGSFVQHPELTGLDCIRPCFWVELIRGRPKENTIWTSGSSI
SFCGVNSDTVGWSWPDGAELPFTIDK*

>N1/19 (A/Wisconsin/588/2019)
*MNTQILVFALIAIIPTNADKI*SAWSHPQFEKGGGGSHHHHHHSSSGSGAHIVMVDAYKPTKGGSGGGG
SIINETADDIVYRLTVIIDDRYESLKNLITLRADRLEMIINDNVSTILAGGSGTGSVKLAGNSSLCPV
SGWAIYSKDNSVRIGSKGDVFVIREPFISCSPLECRTFFLTQGALLNDKHSNGTIKDRSPYRTLMSCP
IGEVPSPYNSRFESVAWSASACHDGTNWLTIGISGPDSGAVAVLKYNGIITDTIKSWRNKILRTQESE
CACVNGSCFTIMTDGPSDGQASYKIFRIEKGKIIKSVEMKAPNYHYEECSCYPDSSEITCVCRDNWHG
SNRPWVSFNQNLEYQMGYICSGVFGDNPRPNDKTGSCGPVSSNGANGVKGFSFKYGNGVWIGRTKSIS
SRKGFEMIWDPNGWTGTDNKFSKKQDIVGINEWSGYSGSFVQHPELTGLNCIRPCFWVELIRGRPEEN
TIWTSGSSISFCGVDSDIVGWSWPDGAELPFTIDK*

>N1/09 hybrid (N1/09 Loops - mSN1 Scaffold)
*MNTQILVFALIAIIPTNADKI*SAWSHPQFEKGGGGSHHHHHHSSSGSGAHIVMVDAYKPTKGGSGGGG
SIINETADDIVYRLTVIIDDRYESLKNLITLRADRLEMIINDNVSTILAGGSGTGSVTLAGNSSLCPI
SGWAIYSKDNGIRIGSKGDVFVIREPFISCSHLECRTFFLTQGALLNDKHSNGTIKDRSPYRTLMSCP
VGEAPSPYNSRFESVAWSASACHDGISWLTIGISGPDNGAVAVVKYNGIITDTIKSWRNNILRTQESE
CACVNGSCFTVMTDGPSNGQASYKIFKIEKGKVVKSVEMNAPNYHYEECSCYPDAGDIMCVCRDNWHG
SNRPWVSFNQNLEYQIGYICSGIFGDNPRPNDKTGSCGPVSSNGANGVKGFSFKYGNGVWIGRTKSIS
SRNGFEMIWDPNGWTETDSSFSVKQDIVGINEWSGYSGSFVQHPEMTGLDCMRPCFWVELIRGRPKEN
TIWTSGSSISFCGVNSDTVGWSWPDGAELPFTIDK*

>N1/19 hybrid (N1/19 Loops - mSN1 Scaffold)
*MNTQILVFALIAIIPTNADKI*SAWSHPQFEKGGGGSHHHHHHSSSGSGAHIVMVDAYKPTKGGSGGGG
SIINETADDIVYRLTVIIDDRYESLKNLITLRADRLEMIINDNVSTILAGGSGTGSVTLAGNSSLCPI

SGWAIYSKDNGIRIGSKGDVFVIREPFISCSHLECRTFFLTQGALLNDKHSNGTIKDRSPYRTLMSCP
VGEAPSPYNSRFESVAWSASACHDGISWLTIGISGPDNGAVAVVKYNGIITDTIKSWRNKILRTQESE
CACVNGSCFTVMTDGPSDGQASYKIFKIEKGKVVKSVEMKAPNYHYEECSCYPDAGDIMCVCRDNWHG
SNRPWVSFNQNLEYQMGYICSGVFGDNPRPNDKTGSCGPVSSNGANGVKGFSFKYGNGVWIGRTKSIS
SRKGFEMIWDPNGWTETDSSFSVKQDIVGINEWSGYSGSFVQHPEMTGLDCMRPCFWVELIRGRPEEN
TIWTSGSSISFCGVNSDTVGWSWPDGAELPFTIDK*

>PR8 N1 (A/PR/8/1934)
*MNTQILVFALIAIIPTNADKI*SAWSHPQFEKGGGGSHHHHHHSSSGSGAHIVMVDAYKPTKGGSGGGG
SIINETADDIVYRLTVIIDDRYESLKNLITLRADRLEMIINDNVSTILAGGSGTGSVILTGNSSLCPI
RGWAIYSKDNSIRIGSKGDVFVIREPFISCSHLECRTFFLTQGALLNDRHSNGTVKDRSPYRALMSCP
VGEAPSPYNSRFESVAWSASACHDGMGWLTIGISGPDNGAVAVLKYNGIITETIKSWRKKILRTQESE
CACVNGSCFTIMTDGPSDGLASYKIFKIEKGKVTKSIELNAPNSHYEECSCYPDTGKVMCVCRDNWHG
SNRPWVSFDQNLDYQIGYICSGVFGDNPRPKDGTGSCGPVYVDGANGVKGFSYRYGNGVWIGRTKSHS
SRHGFEMIWDPNGWTETDSKFSVRQDVVAMTDWSGYSGSFVQHPELTGLDCIRPCFWVELIRGRPKEK
TIWTSASSISFCGVNSDTVDWSWPDGAELPFTIDK*

>mS hybrid (mSN1 Loops - PR8 Scaffold)
*MNTQILVFALIAIIPTNADKI*SAWSHPQFEKGGGGSHHHHHHSSSGSGAHIVMVDAYKPTKGGSGGGG
SIINETADDIVYRLTVIIDDRYESLKNLITLRADRLEMIINDNVSTILAGGSGTGSVILTGNSSLCPI
RGWAIYSKDNSIRIGSKGDVFVIREPFISCSHLECRTFFLTQGALLNDKHSNGTVKDRSPYRALMSCP
VGEAPSPYNSRFESVAWSASACHDGMGWLTIGISGPDNGAVAVLKYNGIITETIKSWRNNILRTQESE
CACVNGSCFTIMTDGPSNGQASYKIFKIEKGKVTKSIELNAPNYHYEECSCYPDTGKVMCVCRDNWHG
SNRPWVSFDQNLDYQIGYICSGVFGDNPRPNDGTGSCSPMSSNGAYGVKGFSYRYGNGVWIGRTKSTS
SRSGFEMIWDPNGWTETDSKFSVRQDVVEITDWSGYSGSFVQHPELTGLDCIRPCFWVELIRGRPKEN
TIWTSASSISFCGVNSDTVDWSWPDGAELPFTIDK*

>PR8 hybrid (PR8 N1 Loops - mSN1 Scaffold)
*MNTQILVFALIAIIPTNADKI*SAWSHPQFEKGGGGSHHHHHHSSSGSGAHIVMVDAYKPTKGGSGGGG
SIINETADDIVYRLTVIIDDRYESLKNLITLRADRLEMIINDNVSTILAGGSGTGSVTLAGNSSLCPI
SGWAIYSKDNGIRIGSKGDVFVIREPFISCSHLECRTFFLTQGALLNDRHSNGTVKDRSPYRTLMSCP
VGEAPSPYNSRFESVAWSASACHDGISWLTIGISGPDNGAVAVVKYNGIITDTIKSWRKKILRTQESE
CACVNGSCFTVMTDGPSDGLASYKIFKIEKGKVVKSVELNAPNSHYEECSCYPDAGDIMCVCRDNWHG
SNRPWVSFNQNLEYQIGYICSGVFGDNPRPKDGTGSCGPVYVDGANGVKGFSFKYGNGVWIGRTKSHS
SRHGFEMIWDPNGWTETDSSFSVKQDIVAMTDWSGYSGSFVQHPEMTGLDCMRPCFWVELIRGRPKEK
TIWTSGSSISFCGVNSDTVGWSWPDGAELPFTIDK*
